# Antibody cross-reactivity accounts for widespread appearance of m^1^A in 5’ UTRs

**DOI:** 10.1101/648345

**Authors:** Anya V. Grozhik, Anthony O. Olarerin-George, Miriam Sindelar, Xing Li, Steven S. Gross, Samie R. Jaffrey

## Abstract

*N*^1^-methyladenosine (m^1^A) was recently identified as a new mRNA modification based on its mapping to the 5’ UTRs of thousands of mRNAs with an m^1^A-binding antibody. More recent studies have confirmed the prevalence of m^1^A, while others have questioned it. To address this discrepancy, we mapped m^1^A using ultra-deep RNA-Seq datasets based on m^1^A-induced misincorporations during reverse transcription. Using this approach, we find m^1^A only in the mitochondrial *MT-ND5* transcript. In contrast, when we mapped m^1^A antibody-binding sites at single-nucleotide resolution, we found binding to transcription start nucleotides in mRNA 5’ UTRs. Using different biochemical assays, we find that m^1^A is not present at these sites. Instead, we find that the m^1^A antibody exhibits m^1^A-independent binding to mRNA cap structures. We also tested a new and independently derived m^1^A antibody. We show that this m^1^A antibody lacks m^7^G cap-binding cross-reactivity, and notably does not map to 5’ UTRs in the transcriptome. Our data demonstrate that high-stoichiometry m^1^A sites are rare in the transcriptome and that previous mapping of m^1^A to mRNA 5’ UTRs are due to unintended binding of the m^1^A antibody to m^7^G cap structure in mRNA.

## Introduction

An emerging concept in gene expression regulation is that the fate and function of mRNAs is controlled by dynamic, and in some cases, reversible nucleotide modifications. The initial concept of the epitranscriptome came with the transcriptome-wide mapping of thousands of internally located modified nucleotide *N*^6^-methyladenosine (m^6^A) residues in the transcriptome^1,2^. m^6^A mapping revealed diverse patterns of methylation in thousands of transcripts in various cells and tissues^1,2^.

A major goal in the field of epitranscriptomics has been to establish whether other modified nucleotides are also highly prevalent in mRNA. Two studies in 2016 identified *N*^1^-methyladenosine (m^1^A) as an abundant modification^3,4^. In these studies, m^1^A was mapped by sequencing mRNA fragments immunoprecipitated with a commercial m^1^A antibody from MBL. Hundreds of m^1^A sites were identified in hundreds of mRNAs. One study estimated the average stoichiometry of m^1^A at 20%^3^. Furthermore, most m^1^A sites were located near start codons, which was suggested to enhance translation^3^. Hence, these studies identified m^1^A as an abundant and functional modification affecting hundreds of mRNAs.

Two subsequent studies proposed different maps of m^1^A. One study argued that m^1^A was exceptionally rare in mRNA^5^. In this study, the antibody-bound RNA was reverse transcribed with a reverse transcriptase that efficiently introduces misincorporations at m^1^A. Using this method, the authors found that m^1^A was rarely present in the RNA immunoprecipitated with the m^1^A antibodies^5^. Although mRNA fragments from 5’ UTRs and start codon-proximal regions were immunoprecipitated, these fragments did not generate misincorporations. Thus, the authors proposed that mRNA fragments from the 5’ UTR may be nonspecifically enriched during immunoprecipitation^5^. Only two mRNAs contained high confidence m^1^A sites: *C9orf100* and *MT-ND5*, a cytosolic and a mitochondrial mRNA, respectively^5^. Twelve other mRNAs were also detected at very low stoichiometry.

The second study mapped m^1^A to 740 sites, 473 of which were in mRNA and lncRNA. In the mRNAs, the majority of sites were found in the 5’ UTR; twenty-two of which the authors localized to the first nucleotide of the transcript. Based on their location, the authors proposed that m^1^A forms a novel type of cap structure in which m^1^A immediately follows the 7-methylguanosine (m^7^G) cap of mRNA (m^7^G-ppp-m^1^A). A re-analysis of this data has argued that the few start codon-proximal m^1^A sites mapped in this study were not mapped to the genome properly, and in fact represent transcription start sites^6^.

In summary, very divergent m^1^A maps have been generated. The major question is whether m^1^A sites are indeed at transcription start sites or start codons, or neither, and why these sites are so prominent in m^1^A-mapping studies. Additionally, it is not clear if m^1^A sites are indeed high stoichiometry modifications as reported in the initial study^3^, or low-stoichiometry and rare as reported in another study^5^.

To resolve the question of the prevalence and location of m^1^A in the transcriptome, we used both a high-resolution m^1^A-mapping method as well as a bioinformatic approach, termed misincorporation mapping, to localize m^1^A in the transcriptome. Misincorporation mapping takes advantage of the ability of m^1^A and certain other modified nucleotides to induce misincorporations during the reverse transcription step in RNA-Seq. By using ultra-deep RNA-Seq datasets, we discovered that very few transcriptomic sites generate misincorporations in multiple independent RNA-Seq datasets. Only the *MT-ND5* mitochondrial transcript and the *MALAT1* noncoding RNA generated statistically significant misincorporations, demonstrating the rarity of high stoichiometry m^1^A sites. To understand why misincorporation mapping reveals few m^1^A sites while m^1^A antibodies detect large number of sites, we mapped m^1^A at high resolution using the same MBL antibody used in all previous studies. Our mapping recapitulated the selective binding of the MBL m^1^A antibody to transcription start nucleotides in mRNA. However, we found that the m^1^A antibody also directly binds the m^7^G cap structure, and this m^1^A-independent binding property explains why previous maps showed m^1^A in mRNA 5’ UTR regions. To confirm this, we show that a newly developed m^1^A antibody, which we show does not bind the m^7^G cap, produces a m^1^A map that no longer shows binding to the 5’ end of mRNAs. Overall, our data show that m^1^A and other hard stop nucleotides are rare in mRNA, and demonstrate that antibody cross-reactivity with 5’ caps accounts for the apparent localization of m^1^A to transcription start nucleotides and start codons.

### Misincorporation mapping: transcriptome-wide detection of modified nucleotides from ultra-deep RNA-seq

Given the inconsistency in the different antibody-dependent m^1^A-mapping methods (**Extended Data Fig. 1a-c**), we sought to use an antibody-independent approach to detect m^1^A at single-nucleotide resolution in mRNA. For this approach, we took advantage of ultra-deep RNA-seq datasets and the fact that m^1^A is a “hard stop” modification, meaning it typically arrests cDNA synthesized by standard reverse transcriptases^7,8^ (**Extended Data Fig. 1d**). However, SuperScript III will read through m^1^A at low frequency, resulting in misincorporations that are variable and sequence dependent^7,8^. Most m^1^A-induced misincorporations are A→T transitions that can be detected by sequencing the cDNA^7,8^. This approach can also detect other hard stop modifications, such as 3-methylcytidine (m^3^C), 3-methyluridine (m^3^U), *N*^2^,*N*^2^-dimethylguanosine, and 1-methylguanosine, since these also produce misincorporations^7^. Therefore, the presence of misincorporated nucleotides can directly localize the m^1^A and other hard stop nucleotides in sequencing data^7,8^.

Misincorporations are difficult to distinguish from sequencing errors using standard next-generation sequencing for two major reasons. First, substantial read depth is required to detect m^1^A since m^1^A typically induces a misincorporation in only approximately 20-30% of the cDNAs generated by SuperScript III^7,8^. The misincorporations would thus be particularly difficult to detect for m^1^A residues that exist at low stoichiometries. Second, misincorporations cannot be readily distinguished from stochastic errors originating during PCR amplification of the library or during sequencing. Thus, m^1^A cannot be definitively identified in standard RNA-seq experiments.

To overcome these problems, we developed a bioinformatic approach similar to high-throughput annotation of modified ribonucleotides (HAMR)^9^ to distinguish modified nucleotides from sequencing errors (**Fig. 1a**). In our approach, we used an ultra-deep RNA-seq dataset from blood mononucleocytes comprising approximately 3 billion reads derived from 20 independent sequencing experiments (“replicates”)^10^. The replicates were derived from a single human donor whose genome was sequenced, allowing any differences between the cDNA and genome to be readily detected. As with most RNA-seq datasets, the library does not contain the exact site where the reverse transcriptase terminatates^10^ (see **Extended Data Fig. 1a**). Thus, this dataset cannot be used to map modification-induced reverse transcriptase stop sites. Instead, we localized hard stop modifications by identifying all nucleotide positions that showed misincorporations across multiple replicates (see **Methods**). Notably, for m^1^A, we considered A→T transitions that were present alongside other rare transition types induced by m^1^A^7,8^ (see **Methods**). Importantly, this method only reveals misincorporations, not the identity of the modification; the identity would have to be determined by biochemical methods.

**Figure 1.**
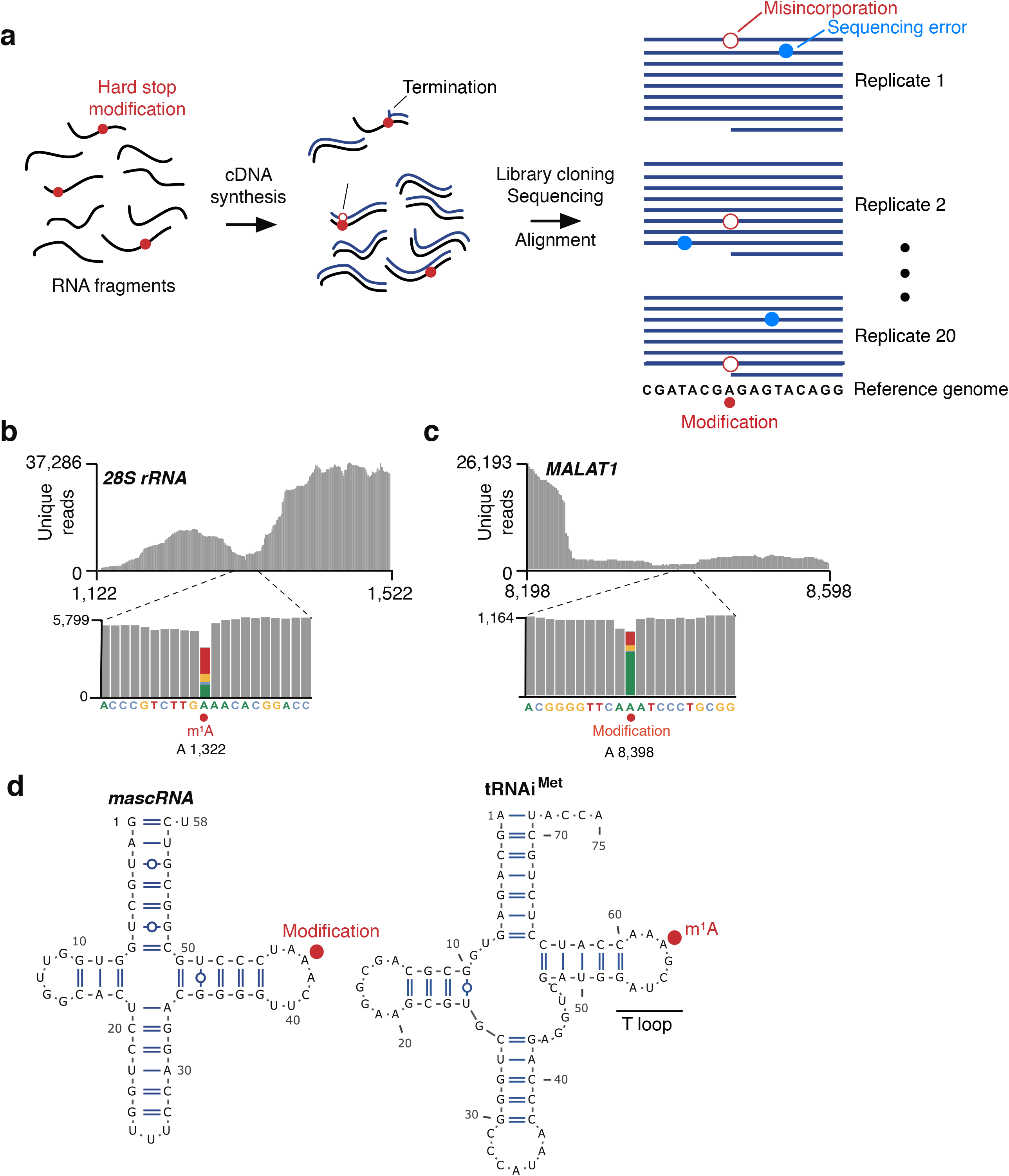
Misincorporation mapping identifies known and novel modifications. **a**, Schematic of misincorporation mapping. An ultra-deep RNA-seq dataset was derived from 20 independent biological replicates. Conceivably, some of the RNA fragments (black lines) contain “hard stop modifications” (red circles), and RNA fragments were reverse transcribed (cDNA, blue lines). For RNA fragments containing these modifications, the reverse transcriptase could terminate at the modification, produce a nucleotide misincorporation, or read through the modification. When the cDNA library is PCR-amplified and sequenced, nucleotide differences are detected between the aligned reads and genomic sequence matching the RNA-seq dataset. Misincorporations (open red circles) identified using this approach are indicative of putative modification sites. Examination of these misincorporations across multiple replicates is critical to distinguish modification sites and sequencing error, which occurs more dispersedly (blue circles). **b**, m^1^A in the *28S* rRNA is detected by misincorporation mapping. To determine if known hard stop nucleotide modifications are detected using misincorporation mapping in ultra-deep RNA-seq, we evaluated mapped sequence reads (grey) around a known m^1^A site in the *28S* rRNA (upper panel). Approximately 70% of the nucleotides that mapped to the m^1^A position contained misincorporations (lower panel; colored bar, position of modified nucleotide and corresponding misincorporations). Additionally, most misincorporations were A→T transitions, typical of m^1^A. **c**, Misincorporation mapping detects a modified adenosine in *MALAT1*. To identify novel modified sites, we analyzed misincorporations in lncRNA and mRNA. A high-confidence site was identified in the lncRNA *MALAT1*. More than 25% of reads (grey) mapping to an adenosine in this RNA contained misincorporations (colored bar, position of modified nucleotide and corresponding misincorporations). **d**, The modification in *MALAT1* described in **c** occurs in tRNA-like structure at a position that corresponds to the m^1^A position in tRNAs. This region of *MALAT1* is known to be processed to yield a tRNA-like small RNA called *mascRNA*^14^ (left), which is similar to human tRNAs (right, tRNAi^Met^ is shown as an example). The location of the modified adenosine in *mascRNA* is analogous to the position of m^1^A in the T-loop structure of tRNA. This result suggests that the modified adenosine in *mascRNA* is likely to contain m^1^A.

We first confirmed that we could detect known m^1^A sites. After aligning reads to rRNA, we readily detected the known m^1^A at position 1,322 in the *28S* rRNA (**Fig. 1b**, **Extended Data Fig. 2a**). As expected, the misincorporations were predominantly A→T transitions, which are characteristic of m^1^A^7,8^. These site-specific misincorporations were detected in all 20 replicates, confirming that the A→T transitions were not stochastic sequencing errors.

Misincorporation mapping can also detect other hard stop modifications. These included 1-methyl-3-(3-amino-3-carboxypropyl)pseudouridine and m^3^U (**Extended Data Fig. 2b**). On the other hand, modifications that do not significantly affect reverse transcription fidelity, such as m^6^A, pseudouridine, *N*^4^-acetylcytidine, 2’-*O*-methylated nucleotides, and m^7^G, did not induce misincorporations (**Extended Data Fig. 2c**). These data demonstrate that hard stop modifications could be reliably detected in this dataset.

We considered the possibility that m^1^A detection could be impaired in this approach, since m^1^A can convert to m^6^A through the Dimroth rearrangement, a heat and base-catalyzed reaction^11^ (**Extended Data Fig. 2d**). To estimate m^1^A loss during the preparation of the ultra-deep sequencing libraries, we examined the m^1^A at position 1,322 in the *28S* rRNA. This site is reported to be methylated at near complete stoichiometry^12^. Since reverse transcription of m^1^A results in read-through approximately 20-30% of the time^7,8^, the fraction of read-through events can suggest the overall stoichiometry of m^1^A. Notably, we found that m^1^A at this position was associated with a read-through rate of approximately 15% in this dataset (see **Extended Data Fig. 2a**). This suggests that the library preparation protocol did not cause substantial depletion of m^1^A, and m^1^A residues should be detectable throughout the transcriptome using this dataset.

### m^1^A-induced misincorporations are not readily detected in mRNA

Next, we wanted to map m^1^A at single-nucleotide resolution in mRNA. Although the number of reads is exceptionally high in this ultra-deep RNA-seq dataset, we reasoned that the read depth may not be sufficient on individual transcripts to detect m^1^A in low abundance mRNAs or mRNAs with low stoichiometry m^1^A. In order to detect m^1^A, an m^1^A residue needs to have been reverse transcribed a sufficient number of times during library preparation to detect misincorporations. Since m^1^A sites were reported to have on average a 20% stoichiometry^3^, we thus set a threshold number of 500 unique reads on any given nucleotide to detect m^1^A sites. At this stoichiometry, 100 reverse transcription events would encounter m^1^A. Of these 100 reverse transcription events, approximately 20% would read through, and most of these would be associated with a misincorporation^7,8^. At this read depth, misincorporations should be readily detected in multiple replicates. Thus, to detect m^1^A in mRNA, we restricted our search to approximately 8 million adenosine residues in the transcriptome that showed a read depth of >500 reads (**Extended Data Fig. 3a**).

Analysis of the 3 billion reads provided by this dataset showed 14 high-confidence nucleotide positions across the transcriptome that showed misincorporations in more than one replicate (see **Methods**). Of these, 12 occurred at adenosine residues (**Extended Data Tables 1,2**). Most of these modified adenosines were found in mitochondrial tRNAs and could be identified as m^1^A since they occurred at known m^1^A positions in mitochondrial tRNAs^13^ (**Extended Data Table 2**). We also detected a modified adenosine in *mascRNA* (*MALAT1*-associated small cytoplasmic RNA), a short tRNA-like ncRNA that is derived from endonucleolytic processing of *MALAT1*^14^ (**Fig. 1c**). Notably, this modified adenosine is found at a position corresponding to position 58 in the T-loop of tRNAs (**Fig. 1d**), a conserved position of m^1^A in tRNAs^15^. This site may be a m^1^A formed by T-loop-specific m^1^A-synthesizing enzymes^15^.

Besides these noncoding RNAs, the previously reported m^1^A-containing *MT-ND5* mitochondrial mRNA contained a modified adenosine (**Extended Data Table 2**). At this modified adenosine, a misincorporation rate of 13.5% was present, in agreement with a recent finding of m^1^A-induced misincorporations at this position in this transcript^5^. This mRNA was also the only mRNA previously found to have relatively high m^1^A stoichiometry (exceeding 50% in a number of cell types and tissues) in a recent reanalysis of m^1^A in the transcriptome^5^. The high degree of misincorporations compared to background levels supports the high sensitivity of this method.

We next examined misincorporations at other mitochondrial mRNAs with annotated m^1^A sites. Studies by Safra *et al*.^5^ and Li *et al*.^16^ identified 11 and 5 putative mitochondrial m^1^A-containing protein-coding genes, respectively. Four mitochondrial mRNAs were common to both studies (*MT-ND5*, *MT-CO1*, *MT-CO2* and *MT-CO3*). Except for *MT-ND5*, the misincorporation rates for these mRNAs were found to be very low when examining purified poly(A) RNA (less than 0.7% and 2.1% in the Safra and Li studies, respectively). However, when these authors enriched m^1^A-containing mRNA using the m^1^A antibody, misincorporations could be detected, suggesting that rare m^1^A-containing transcripts are found in cells. We therefore further examined these sites in our study leveraging the exceptional read depth across the mitochondria (average ~1 million reads/nucleotide). Similarly, we found that with the exception of *MT-ND5* which had a misincorporation rate of 13.5%, the misincorporation rates for all other putative m^1^A sites was less than 0.4% (**Extended Data Fig. 3b, Extended Data Table 3**). Taken together, these earlier studies and our data demonstrate that *MT-ND5* is unique in containing readily detectable m^1^A, and other mitochondrial mRNAs may contain m^1^A, but only at very low stoichiometries.

We also detected three additional sites of modifications in cytosolic mRNAs, with only one site being a modified adenosine (**Extended Data Table 2**). Notably, when we examined cytosolic mRNAs that were previously reported by earlier mapping studies^3^ to have the highest stoichiometry of m^1^A (i.e. >50%) such as *CCDC71*, *DLST*, and *STK16* lacked evidence of A→T transitions based on our misincorporation mapping (**Extended Data Fig. 3c**, **Extended Data Table 1**).

We did not detect any high-confidence misincorporations at cytidine residues in mRNA (**Extended Data Table 2**), in contrast to a recent study that detected m^3^C, a hard-stop nucleotide, in mammalian mRNA by mass spectrometry^17^.

Although misincorporation mapping demonstrated that few mRNAs have m^1^A or other hard stop modifications, we could not conclusively rule out modifications at the very 5’ end of RNAs. This is because RNA-seq typically does not provide coverage at the extreme 5’ ends of RNAs and so detecting misincorporations in this region is not possible (see **Extended Data Fig. 1a**)^18^.

Overall, the paucity of m^1^A sites in mRNA using misincorporation mapping supports the idea that m^1^A is not a highly prevalent and high stoichiometry modification, and is instead exceptionally rare in mRNA.

### m^1^A miCLIP with the MBL m^1^A antibody detects known m^1^A residues at nucleotide resolution

Since misincorporation mapping did not show evidence for m^1^A sites in the vicinity of start codons, we wanted to understand the source of this 5’ UTR signal in the antibody-based mapping approaches. We therefore developed an approach to detect m^1^A at single-nucleotide resolution, m^1^A miCLIP (m^1^A modified individual nucleotide resolution crosslinking and immunoprecipitation) (**Fig. 2a**). In this method, we used the MBL m^1^A antibody that was previously characterized and used to map m^1^A^3,4^. However, m^1^A miCLIP takes advantage of m^1^A antibody-induced crosslinks in mRNA to efficiently terminate cDNA synthesis. Thus, the sites of reverse transcription terminations act as a signature to localize precise sites where the m^1^A antibody binds in mRNA.

**Figure 2.**
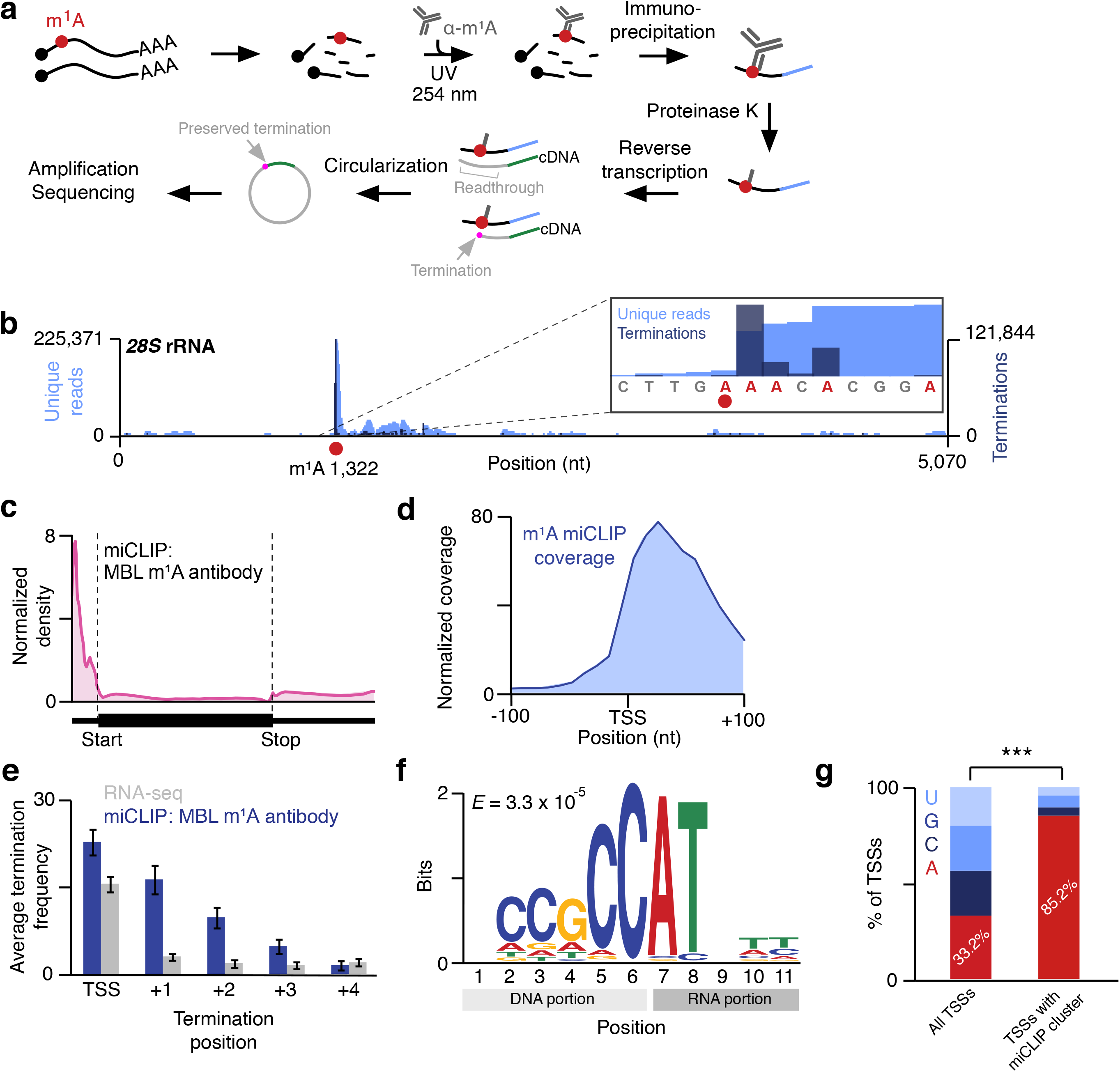
Mapping m^1^A in the transcriptome using m^1^A miCLIP. **a**, m^1^A miCLIP workflow. For transcriptome-wide m^1^A (red circles) profiling at nucleotide resolution, poly(A) RNA (black lines; black circles, RNA caps) was fragmented, incubated with m^1^A antibody, and crosslinked to the antibody with 254 nm UV light. RNA-antibody complexes were immunoprecipitated, ligated to a 3’ linker (blue line), and purified. RNA was released from crosslinked antibody using Proteinase K, generating RNA fragments containing crosslinked peptide. RNA fragments were reverse transcribed, and the first-strand cDNA (grey lines) was circularized to preserve potential m^1^A-induced read terminations (magenta dot) at the cDNA 3’ end. Circularized cDNA was then subjected to library construction. Amplified libraries were subjected to Illumina sequencing. **b**, m^1^A miCLIP detects the known m^1^A site in the *28S* rRNA. m^1^A miCLIP unique reads (light blue) were aligned to rRNA, and the m^1^A site (red circle) at position 1,322 of the *28S* rRNA was examined. m^1^A miCLIP reads were strikingly enriched at the m^1^A site but not at other locations along the *28S* rRNA. Moreover, the reads terminated predominantly at the +1 position relative to the m^1^A (inset, dark blue), consistent with the terminating effect of m^1^A on reverse transcription. This demonstrates that m^1^A miCLIP detects m^1^A sites with high specificity and resolution. **c**, Relative distribution of m^1^A miCLIP clusters in mRNA. The metagene distribution of m^1^A miCLIP clusters that were found in mRNA and contained 20 or more stacked reads (*N* = 3,877) was normalized to RNA-seq coverage and plotted. Clusters were predominantly enriched in the 5’ UTR and next to the transcription start site (start, start codon; stop, stop codon). The annotation of transcription start sites is not exact, which results in slight coverage upstream of the transcription start site. **d**, m^1^A miCLIP coverage is enriched at and near transcription start sites. m^1^A miCLIP unique reads were mapped relative to annotated transcription start sites using Deeptools (see **Methods**). **e**, The m^1^A antibody produces crosslink-induced terminations of reverse transcription near the transcription start site. The termination sites of m^1^A miCLIP reads (blue) and RNA-seq reads (grey) are shown relative to the transcription start site. While RNA-seq reads terminate predominantly at position 0 (i.e. due to the end of the transcript), m^1^A miCLIP reads terminate predominantly at positions 0 and +1, and to a lesser degree, at downstream nucleotides, suggesting that the m^1^A antibody binds at or near the transcription start site and blocks reverse transcription. **f**, The transcription initiation motif is enriched by m^1^A miCLIP. m^1^A miCLIP clusters found in 5’ UTRs were annotated with their genomic sequence environment, and the sequences were analyzed for recurrent motifs. This resulted in significant enrichment of a pyrimidine-rich sequence motif that facilitates transcription initiation of mRNAs at adenosine (*E* = 3.3 × 10^−5^). Genomic and transcribed regions of the sequence motif are marked. **g**, Adenosine transcription start sites are enriched in m^1^A miCLIP clusters. m^1^A miCLIP clusters were overlapped with a collective set of transcription start sites (TSS) that included RefSeq annotations and recently mapped high-resolution regions containing m^6^Am mRNA extended caps^43^, a form of initiating adenosine in certain mRNAs. While 33.2% of these collective transcription start sites were adenosines, the frequency of adenosine transcription start sites overlapping m^1^A miCLIP clusters was significantly higher (85.2%, *N* = 1,026 TSSs, *** p < 2.2 × 10^−16^, Fisher’s exact test).

In m^1^A miCLIP, we crosslink the m^1^A-specific antibody to sheared RNA (**Fig. 2a**). UV crosslinking was previously shown to reduce nonspecific RNA binding and increase peak resolution in mapping studies^19^. RNA fragments crosslinked to the antibody were then purified and cloned as a cDNA library. Terminations introduced during reverse transcription were then analyzed to determine the positions of m^1^A throughout the transcriptome.

The library generated in m^1^A miCLIP is different from the initial m^1^A mapping methods^3,4^ since those libraries are generated in way that the cDNA ends are not preserved and produce peaks that are displaced 3’ from the actual location of the m^1^A (see **Extended Data Fig. 1a**). We chose to preserve the cDNA ends since they can identify the exact sites of m^1^A-induced terminations. m^1^A miCLIP is also not affected by peak displacement since it maps the actual site of m^1^A by detecting the precise positions of m^1^A antibody-binding events within transcripts.

Prior to performing transcriptome-wide m^1^A mapping, we confirmed the previously described specificity of the MBL m^1^A antibody. In these experiments, we monitored the ability of competitor nucleotides to prevent the m^1^A antibody from crosslinking to cellular RNA. Only m^1^A abolished antibody-RNA crosslinking (**Extended Data Fig. 4a**), confirming that the antibody binds m^1^A, and not other nucleotides.

Next, we examined the ability of m^1^A miCLIP to detect known m^1^A sites. We performed m^1^A miCLIP with the MBL antibody using poly(A) RNA from HEK 293T cells and examined termination signatures at m^1^A sites in rRNA and tRNA, which typically co-purify to some extent with poly(A) RNA^20-22^. The majority of reads truncated at the +1 position relative to the m^1^A at position 1,322 of the *28S* rRNA (**Fig. 2b**, **Extended Data Fig. 4b**) as well as known m^1^A sites in tRNA (**Extended Data Fig. 4c**). As expected, some read-through was also observed, reflecting the low read-through rate of SuperScript III when it encounters m^1^A. These data demonstrate specific detection of m^1^A by m^1^A miCLIP using the MBL m^1^A antibody.

Notably, m^1^A miCLIP showed markedly improved peak resolution compared to the previous peak-based m^1^A mapping studies^3,4^ (**Extended Data Fig. 4b**).

We asked if m^1^A detection in m^1^A miCLIP is affected by the known instability of m^1^A, potentially causing missed or undetected m^1^A sites. m^1^A converts to m^6^A when exposed to high temperatures and basic pH^11^, which were not used in m^1^A miCLIP (see **Extended Data Fig. 2d**). Nevertheless, to determine if m^1^A converts to m^6^A during m^1^A miCLIP, we asked if m^6^A is detected at m^1^A sites in rRNA. To test this, we analyzed the distribution of m^6^A miCLIP reads on rRNA using a recently-published m^6^A miCLIP dataset^23^. As expected, the known m^6^A site at position 4,220 in *28S* rRNA was detected in m^6^A miCLIP. However, no m^6^A peak above noise level was detected at position 1,322, the m^1^A site in *28S* rRNA (**Extended Data Fig. 4d**). Thus, m^1^A does not appreciably convert to m^6^A and is highly stable during the m^1^A miCLIP protocol.

Taken together, these data indicate that m^1^A miCLIP maps m^1^A with high specificity and resolution.

### The MBL m^1^A antibody binds near the first transcribed nucleotide of mRNA

We next used m^1^A miCLIP to map m^1^A in mRNA (**Extended Data Fig. 5a**). To do this, we aligned m^1^A miCLIP unique reads to the genome, generated m^1^A miCLIP clusters (see **Methods**, **Extended Data Table 4**), and analyzed their distribution. A metagene analysis of m^1^A miCLIP clusters obtained from HEK 293T cells showed a marked enrichment in the 5’ UTR (**Fig. 2c**). More precisely, the clusters were located at mRNA transcription start sites (**Fig. 2d**). A similar enrichment was seen in m^1^A miCLIP clusters obtained from mouse mRNA (**Extended Data Table 5**, **Extended Data Fig. 5b**).

The location of m^1^A antibody crosslinks in mRNA suggested that the MBL m^1^A antibody binds at or near the transcription start nucleotide of diverse mRNAs. To confirm that the MBL m^1^A antibody binds the transcription start nucleotide, we examined the pattern of crosslink-induced terminations around transcription start nucleotide of bound mRNAs. Crosslinking of the m^1^A antibody is expected to cause reverse transcription terminations within several nucleotides of the site of the antibody-RNA adduct^19^. As expected, we found that in miCLIP, terminations were enriched not only at the transcription start site, but also prominently at the +1 position relative to the transcription start site (**Fig. 2e**), with additional terminations sometimes seen between position +2 and +3 (**Fig. 2e**). Thus, the MBL m^1^A antibody binds at or near mRNA transcription start nucleotides.

We considered the possibility that terminations near the transcription start nucleotide could simply reflect general behavior of the reverse transcriptase as it approaches the 5’ end of the mRNA. To test this, we examined the input RNA fragments in the RNA-seq dataset prepared using the same library cloning strategy as m^1^A miCLIP. In general, RNA-seq reads terminated almost exclusively at the transcription start nucleotide, while m^1^A miCLIP reads often terminated at position +1, and up to +3, and in cases of some transcripts, up to +5 (**Fig. 2e**, **Extended Data Fig. 5c**). Therefore, read terminations seen in m^1^A miCLIP near the transcription start nucleotide are likely induced by selective binding and crosslinking of the MBL m^1^A antibody rather than an artifact of reverse transcription near mRNA 5’ ends.

Since m^1^A miCLIP clusters predominantly localize to the 5’ UTR, we analyzed clusters found in this region for the presence of any consensus sequence. This analysis revealed a consensus sequence that was pyrimidine-rich (**Fig. 2f**) and highly similar to Initiator, a transcription-initiating sequence which produces transcripts that initiate with adenosine^24,25^. This finding further supports that the MBL m^1^A antibody likely binds at or near the transcription start nucleotide.

We further determined that 85% of transcripts containing an m^1^A miCLIP cluster at their transcription start nucleotide initiated with adenosine (**Fig. 2g**, **Extended Data Table 6**). Thus, we reasoned that adenosine at the transcription start nucleotide was important for binding of the antibody at or near the transcription start site.

### m^1^A and m^1^A_m_ are not detected in mRNA extended cap structures

Because the MBL m^1^A antibody binds at transcription start sites, it has been proposed that mRNAs bound by this antibody contain mRNA cap structures containing m^7^G followed by *N*^1^-methylated adenine at the transcription start nucleotide^16^. The methylated nucleotide would be m^1^A or *N*^1^,2’-*O*-dimethyladenosine (m^1^A_m_). m^1^A_m_ may be more likely since the first encoded nucleotide of mRNAs is typically subjected to 2’-*O*-methylation^26,27^. To biochemically validate this hypothesis, we first used mass spectrometry to detect m^1^A or m^1^A_m_ within extended cap structures. We specifically sought to detect m^1^A or m^1^A_m_ in the context of the “cap dinucleotide,” i.e., m^7^G-ppp-m^1^A_(m)_. We therefore treated cellular RNA with P1 nuclease, which digests all internal nucleotides to mononucleotides, but leaves the cap dinucleotide intact (see **Methods**).

We readily detected cap dinucleotides using high-resolution liquid chromatography and mass spectrometry using positive ion mode detection. To achieve this, we developed a multiple reaction monitoring protocol based on the fragment ion transitions from distinct dinucleotide precursor species (see **Methods**). To confirm that these *N*^1^-methylated adenosine in a cap dinucleotide can be detected, we used synthetic RNA standards (see **Methods**). Using the standards, we readily detected m^7^G-ppp-m^1^A as well as other cap dinucleotides, such as m^7^G-ppp-A_m_, and m^7^G-ppp-m^6^A_m_ (**Fig. 3a**).

**Figure 3.**
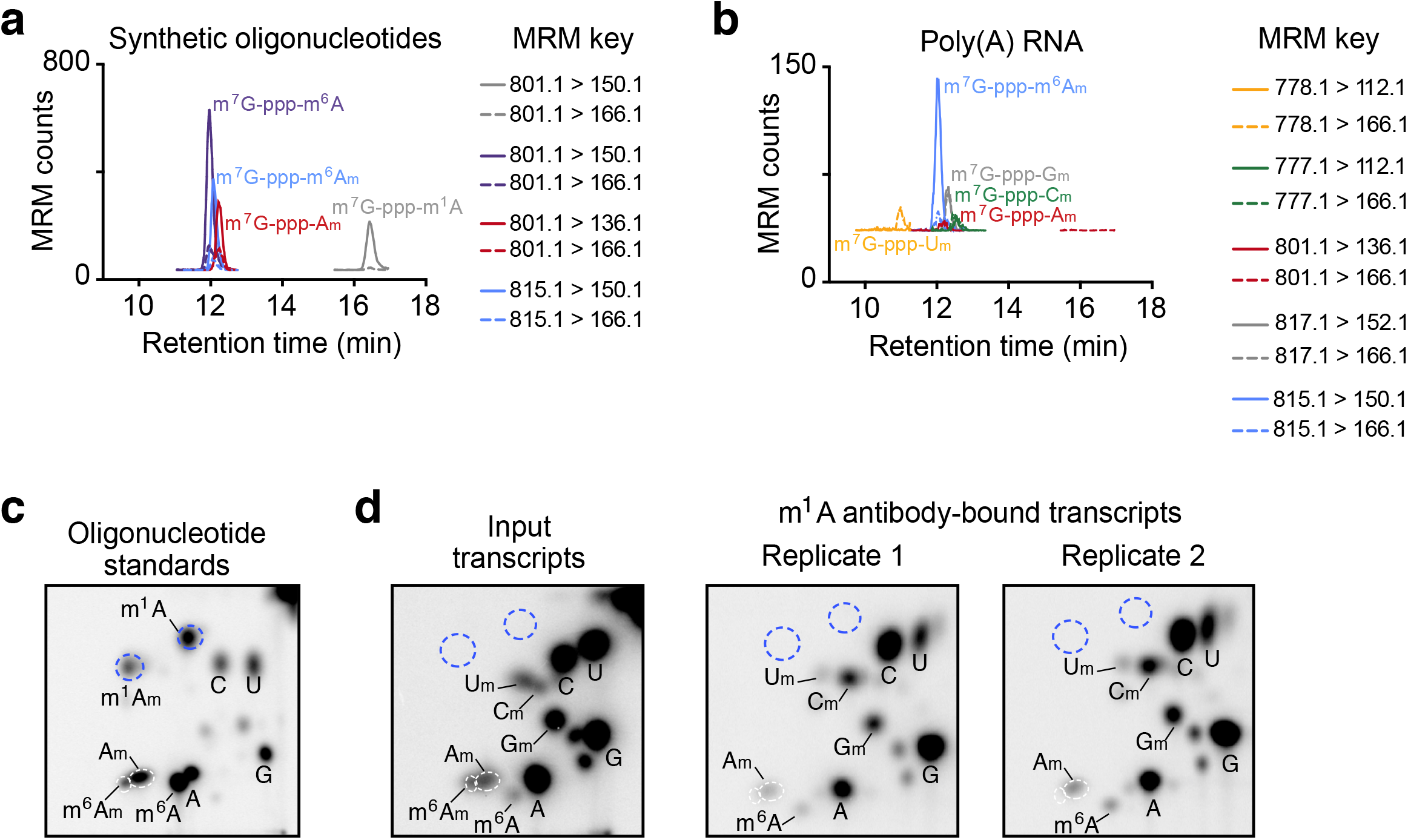
The MBL m^1^A antibody does not enrich for a novel m^1^A-containing cap structure. **a**, Detection of synthetic 5’ cap dinucleotides using mass spectrometry. Dynamic multiple reaction monitoring (dMRM) chromatograms were recorded in positive ion monitoring. Injected was a mixture of oligonucleotides containing m^7^G-ppp-m^6^A_m_, m^7^G-ppp-m^1^A, m^7^G-ppp-m^6^A, or m^7^G-ppp-A_m_ cap structures. Two MRM transitions were recorded for each standard (indicated in key in figure). Note the permanent positive charge on the adenine base of m^7^G-ppp-m^1^A resulted in an increased retention time on the ANP resin as expected. Thus, the m^7^G-ppp-m^1^A_m_ dinucleotide would be expected to elute at a retention time later than m^7^G-ppp-m^6^A due to the permanently positive charged adenine, and earlier than m^7^G-ppp-m^1^A due to the additional methyl group on the ribose. **b**, Detection of 5’ cap dinucleotides from cellular mRNA using mass spectrometry. Dynamic multiple reaction monitoring (dMRM) chromatograms were recorded in positive ion monitoring. Injected were mRNA samples from HEK 293T cells. Two MRM transitions were recorded for each cap structure of interest (indicated in key in figure). We searched for peaks containing MRM transitions and retention times reflecting m^7^G-ppp-m^1^A or m^7^G-ppp-m^1^A_m_ (see note in **a**), but no such signal was detected. **c**, 2D-TLC analysis of capped oligonucleotide standards containing various modified adenines as initiating nucleotides. The migration properties of these various initiating nucleotides are indicated (here, the initiating nucleotides include m^1^A, m^1^A_m_, A_m_, m^6^A, and m^6^A_m_). Spots reflecting A, C, G, and U are also marked. Unmarked spots most likely reflect RNA dinucleotides that are not completely digested during RNA preparation^44^. **d**, 2D-TLC analysis of mRNA extended caps in antibody-unbound (left panel) or m^1^A antibody-bound poly(A) RNA. Known identities of detected spots are marked. Expected positions of potential m^1^A or m^1^A_m_-initiating nucleotides are marked (blue dashed circles) based on standard migrations in **a**.

We next examined cap dinucleotides prepared by digesting poly(A) RNA from HEK293T cells. We readily detected m^7^G-ppp-m^6^A_m_ (*m/z* = 815.1) and also m^7^G-ppp-A_m_ (*m/z* = 801.1), though to a lower degree. Additional species that were detected included extended caps of mRNAs initiating with nucleotides other than adenosine, such as m^7^G-ppp-C_m_ (*m/z* = 777.1), m^7^G-ppp-G_m_ (*m/z* = 817.1), and m^7^G-ppp-U_m_ (*m/z* = 778.1) (**Fig. 3b**). The identity of each species was confirmed by detection of fragment masses corresponding to 7-methylguanine (*m/z* = 166.1) and the base comprising the first nucleotide (*m/z* = m^6^A_m_ 150.1, A_m_ 136.1, C_m_ 112.1, G_m_ 152.1, U_m_ 112.1) within the extended cap.

Next, we examined whether either m^7^G-ppp-m^1^A or m^7^G-ppp-m^1^A_m_ is present in mRNA. Importantly, the masses of these structures (*m/z* = 801.1 and 815.1, respectively) are identical to cap dinucleotides that would contain A_m_ or m^6^A_m_. Moreover, the mass of the fragment produced by the *N*^1^-methylated adenine base (*m/z* = 150.1) would be identical to that produced by the m^6^A_m_ cap. However, m^7^G-ppp-m^1^A and m^7^G-ppp-m^1^A_m_ exhibit distinct retention times from the A_m_ and m^6^A_m_ cap dinucleotides based on our synthetic RNA standards (see **Fig. 3a**), permitting differentiation of *N*^1^-methyl- and *N*^6^-methyl-containing adenine. Nevertheless, we found that no *N*^1^-methylated adenine-containing cap dinucleotide was detected in mRNA (**Fig. 3b**). Thus, m^7^G-ppp-A_m_ or m^7^G-ppp-m^6^A_m_ cap structures were readily detected, while m^7^G-ppp-m^1^A was not detectable.

Since our mass spectrometry analysis does not support the idea that m^1^A was present at the transcription start nucleotide, we wanted to directly and sensitively determine the identity of the transcription start nucleotide that is recovered by the m^1^A antibody. We therefore used two-dimensional thin-layer chromatography (2D-TLC), which can be used to identify and quantify the first encoded nucleotide^28^. In this approach, mRNA is decapped, and the exposed 5’ end of the mRNA is radiolabeled, permitting sensitive detection of the first transcribed nucleotide. The radiolabeled nucleotide species are then resolved using 2D-TLC^28^. Based in the mobility of each species, the identity of the transcription start nucleotide can be determined.

We subjected poly(A) RNA and poly(A) RNA enriched with the m^1^A antibody to 2D-TLC analysis of the transcription start nucleotide. In these experiments, we also used synthetic RNA oligonucleotides containing m^7^G-ppp-m^1^A and m^7^G-ppp-m^1^A_m_ extended cap structures as standards (see **Methods**). Using these standards, we first optimized the solvent conditions so that m^1^A and m^1^A_m_ migrated to distinct positions on the 2D-TLC (**Fig. 3c**). In poly(A) RNA, no m^1^A or m^1^A_m_ was detected at transcription start nucleotides (**Fig. 3d**). Since these might be rare nucleotides at transcription start nucleotides, we enriched for transcription start m^1^A or m^1^A_m_-containing mRNAs by immunoprecipitating poly(A) mRNA with the MBL m^1^A antibody before 2D-TLC. Here, we again did not see m^1^A or m^1^A_m_ as the transcription start nucleotide (**Fig. 3d**).

Taken together, the mass spectrometry and the TLC data suggest that m^1^A and m^1^A_m_ are not readily detectable at the transcription start nucleotide, and m^7^G-ppp-m^1^A_(m)_ does not constitute a novel and prevalent mRNA cap structure as recently proposed^16^.

### The MBL m^1^A antibody directly binds m^7^G-ppp cap structures

Although the antibody-based m^1^A mapping approach suggested that the MBL m^1^A antibody binds at the transcription start nucleotide, we did not observe m^1^A or m^1^A_m_ at this site by either mass spectrometry or TLC. Therefore, we wondered if the MBL antibody binds this region in an m^1^A-independent manner. When we originally characterized the antibody, we performed classic competition studies using nucleosides or nucleotides. However, based on the binding properties of the antibody revealed by mapping studies, we considered the possibility that the MBL m^1^A antibody could bind a larger epitope comprising the mRNA extended cap.

To test this, we wanted to assess whether the MBL m^1^A antibody can bind m^7^G and other modified nucleotides. For binding assays, it is not appropriate to spot m^7^G-containing RNAs since it is not known how m^7^G cap structures interact with membranes, and if its interaction may obscure epitopes needed for m^1^A antibody recognition. We therefore used the standard and widely used competition approach for assessing whether m^7^G extended cap structures bind the MBL m^1^A antibody. We therefore used a dot blot assay in which m^1^A-containing oligonucleotides were spotted on membranes, and the MBL m^1^A antibody was incubated with the membrane in the presence of various competitors. As expected, competition with m^1^A nucleotide inhibited antibody binding (**Fig. 4a**). Related nucleotides, including adenosine, m^6^A, ethenoadenosine, and *N*^1^-substituted nucleotides, like *N*^1,6^-dimethyladenosine (m^1,6^A), did not compete with binding (**Fig. 4a**). However, a commercially available cap analog, m^7^G-ppp-A, was a relatively effective competitor, with an IC_50_ of 480 nM compared to 100 nM for m^1^A (**Fig. 4b**). Thus, the MBL m^1^A antibody can bind the adenine-containing extended cap structure.

**Figure 4.**
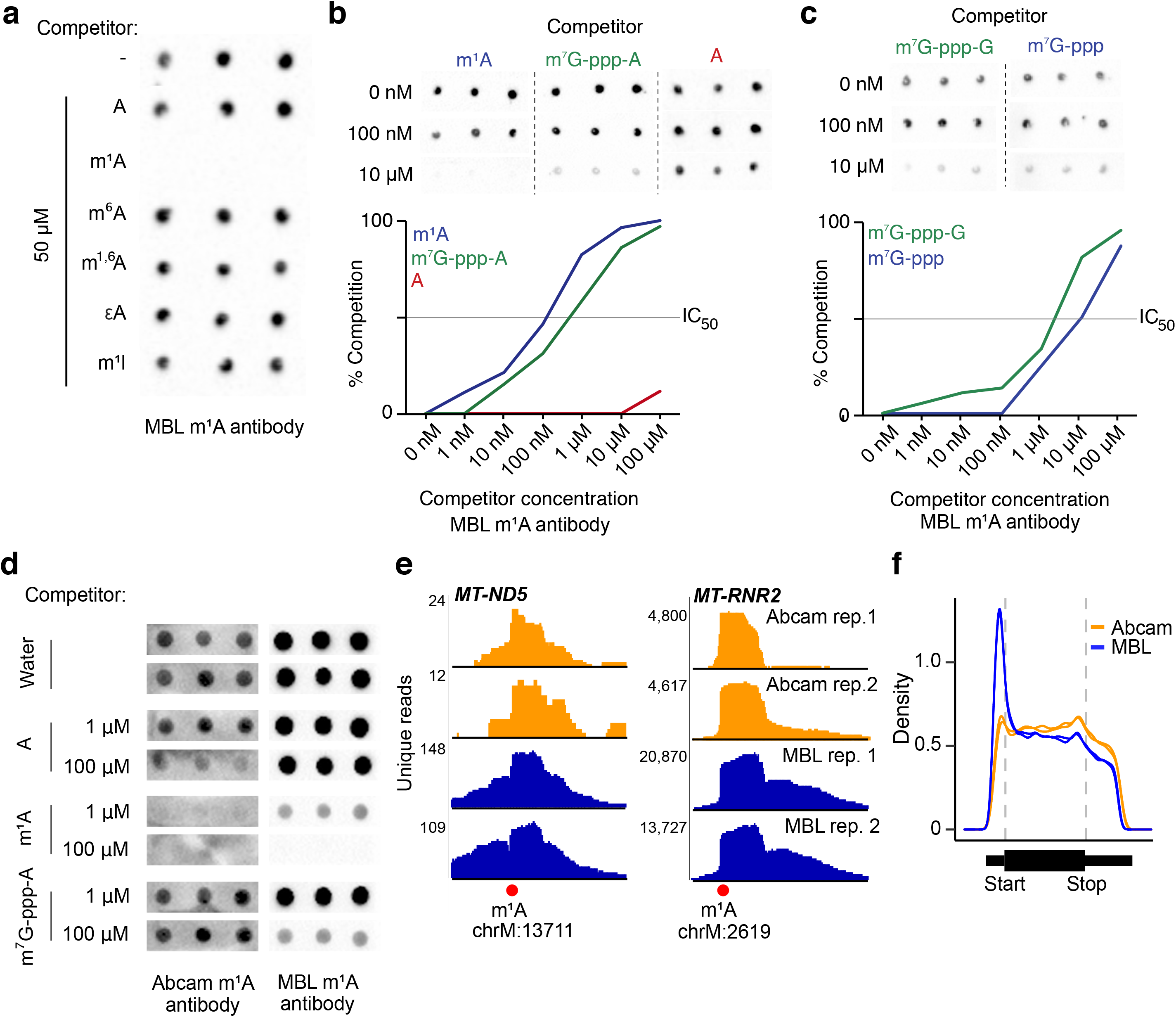
The MBL m^1^A antibody cross-reacts with the mRNA cap structure resulting in enrichment of reads in the 5’ UTR. **a**, The MBL m^1^A antibody is specific for m^1^A compared to other m^1^A-like nucleosides. Shown is a dot blot of an m^1^A-containing oligonucleotide spotted on a nylon membrane in triplicate. The membrane was incubated with the MBL m^1^A antibody along with one of the seven indicated nucleosides similar to m^1^A at the indicated concentration. Only m^1^A was able to prevent the MBL antibody from binding to the spotted oligonucleotide. m^1,6^A, *N*^1,6^-methyladenosine; εA, ethenoadenosine; m^1^I, 1-methylinosine. **b**, Structure-activity relationships between the m^1^A antibody and various substrates. Top, images of dot blot competition assays^1^ where binding of the m^1^A antibody to an m^1^A-containing oligonucleotide was competed with indicated concentrations of m^1^A (m^1^ATP), m^7^G-ppp-A, or A (ATP). Results of three representative competitor concentrations are shown for each competitor. Bottom, quantification of top plus additional concentrations of the competing molecules. Notably, the ability of m^7^G-ppp-A (IC_50_ < 1 μM) to compete with antibody binding is only approximately 10-fold lower than that of the expected specific competitor m^1^A (IC_50_ = ~100 nM). **c**, The MBL m^1^A antibody recognizes the m^7^G-ppp portion of the cap structure. Shown is a dot blot analysis of an m^1^A-containing oligonucleotide spotted on a nylon membrane in triplicate. The membrane was incubated with the MBL m^1^A antibody along with m^7^G-ppp or m^7^G-ppp-G at the indicated concentrations. Both m^7^G-ppp and m^7^G-ppp-G competed with the m^1^A oligonucleotide for binding to the antibody (approximate IC_50_ = 4 μM for m^7^G-ppp-G, 10 μM for m^7^G-ppp). While the antibody’s affinity for m^7^G-ppp-A is higher than for either of these molecules, this result indicates that the m^1^A antibody likely at least partly recognizes extended caps through interactions with the m^7^G-ppp structure. **d**, The Abcam m^1^A antibody does not bind the mRNA cap. Shown are dot blots of an m^1^A-containing oligonucleotide spotted on a nylon membrane in triplicate. The membranes were incubated with either of the two m^1^A antibodies (MBL or Abcam) in the presence of the indicated competing molecules (A, m^1^A or m^7^G-ppp-A; A and m^1^A were either tri- or mono-phosphorylated) at the indicated concentrations. While both antibodies were competed by m^1^A, only the MBL antibody was also competed by m^7^G-ppp-A. Thus, unlike the MBL antibody, the Abcam antibody is selective for m^1^A, relative to the mRNA cap. **e**, The m^1^A antibodies from Abcam and MBL both detect established m^1^A sites. m^1^A miCLIP unique reads from two independent antibodies (in replicate) were aligned to the genome, and the m^1^A sites (red circle) in *MT-ND5* and the mitochondrial 16S rRNA (*MT-RNR2*) were examined. Both antibodies resulted in peaks at the indicated sites with a characteristic truncation right at the m^1^A site. Hence, both m^1^A antibodies specifically recognize known m^1^A sites. **f**, The m^1^A antibody from Abcam lacks the prominent 5’ UTR metagene peak seen with the MBL antibody. As in **e**, unique m^1^A miCLIP reads were aligned to the genome and the relative location of the reads in each transcript is shown with a metagene. Each trace is an independent biological replicate using the indicated antibody. The 5’ UTR peak is essentially absent when the m^1^A-specific Abcam antibody was used compared to the cap-cross-reacting MBL antibody.

We next further characterized the cap-binding properties of the MBL m^1^A antibody. Based on our miCLIP analysis, transcription start site-associated m^1^A read clusters were preferentially enriched on mRNAs that initiated with adenosine (see **Fig. 2g**). Hence, we asked whether the antibody had a preference for adenosine after the m^7^G compared to other nucleotides. To test this, we examined the binding of the MBL m^1^A antibody to the only other commercially available cap analog, m^7^G-ppp-G. The MBL m^1^A antibody bound to m^7^G-ppp-G, but at a weaker IC_50_ (~4 μM) than m^7^G-ppp-A (**Fig. 4c**). The higher binding to the m^7^G-ppp-A cap analog compared to the m^7^G-ppp-G cap analog may explain the preferential binding of the MBL m^1^A antibody to mRNAs that initiate with adenosine. Notably, the antibody also showed binding to m^7^G-ppp, but not m^7^G or ATP, (**Fig. 4c**, **Extended Data Fig. 4a**), demonstrating that the antibody’s binding specificity includes recognition of features all along the entire m^7^G-ppp-A extended cap structure.

Since the MBL m^1^A antibody binds the extended cap structure in an m^1^A-independent manner, we speculated that the m^1^A peaks seen in the 5’ UTR when using this antibody simply reflect binding to the mRNA cap. This would explain why our m^1^A miCLIP shows read enrichment at the transcription start nucleotide, as has also been seen using other m^1^A mapping approaches^16^.

To test this idea, we wanted to examine m^1^A peaks using an antibody that does not show cross-reactivity with the mRNA cap. We therefore tested an additional m^1^A antibody from Abcam that recently became commercially available. Unlike the MBL m^1^A antibody, the Abcam m^1^A antibody was specific for m^1^A but did not show competition with the m^7^G-ppp-A cap analog (**Fig. 4d**). Therefore, the Abcam antibody does not bind the mRNA cap.

We first confirmed that m^1^A miCLIP using HEK293T poly(A) RNA and either the Abcam or MBL antibody detects m^1^A. Indeed, in both cases, we observed a robust peak at the m^1^A sites in *MT-ND5* and *MT-RNR2*, the mitochondrially encoded *16S* RNA (**Fig. 4e**), confirming the ability of each antibody to detect validated m^1^A sites in mRNA.

We next asked if m^1^A miCLIP performed using the m^1^A-specific Abcam antibody showed the same transcriptome-wide enrichment of m^1^A-containing fragments in the 5’ UTR as seen with the MBL m^1^A antibody^3,4^. As expected, the metagene of the miCLIP fragments using the MBL antibody showed a prominent 5’ UTR enrichment (**Fig. 4f**). However, a metagene analysis of all the immunoprecipitated reads using the Abcam m^1^A antibody lacked the 5’ UTR enrichment (**Fig. 4f**). Together, these data demonstrate that only the MBL m^1^A antibody, which cross-reacts with the cap, shows a 5’ UTR enrichment in read coverage. Overall, these data demonstrate that the binding to the 5’ UTR regions is not related to the presence of m^1^A at these sites but is instead a product of antibody cross-reactivity.

### Comparison of m^1^A miCLIP with earlier m^1^A maps

Although one m^1^A mapping study argued that only *MT-ND5* and a few other transcriptomic sites contain m^1^A^5^, other studies showed that both the transcription start nucleotide and start codon-associated 5’ UTR regions contain m^1^A^3,416^. Although the antibody cross-reactivity with mRNA cap structures explains why m^1^A was mapped to transcription start nucleotides, our data does not explain the localization of m^1^A to start codon regions, which was proposed to mediate a novel form of translation initiation^3^. We therefore wanted to understand the exact location of these 5’ UTR- and start codon-associated putative m^1^A sites in light of our misincorporation mapping results, our m^1^A miCLIP using the MBL m^1^A antibody, and our new understanding of the cap-binding properties of the MBL m^1^A antibody.

When we compared mRNAs containing MBL m^1^A antibody miCLIP coverage in this study with the Dominissini *et al.* m^1^A map that localized m^1^A to start codons^3^, we found that the majority of transcripts captured by m^1^A miCLIP were also detected by Dominissini *et al.* (**Extended Data Fig. 6a**). However, the location of reads was different (**Extended Data Fig. 6b**). In particular, the 5’ ends of the miCLIP reads approached the transcription start site, while reads from Dominissini *et al.* were located downstream of the transcription start site (**Extended Data Fig. 6b**, insets). This lateral displacement of peaks towards the start codon is consistent with the library cloning method used in this earlier method (see **Extended Data Fig. 1a**).

In earlier m^1^A mapping studies, accumulations of reads, or in some cases, “troughs” of reduced read coverage due to a putative m^1^A site, were used to predict m^1^A residues to start codons in mRNA^3^. Unlike peaks and troughs, which are a common nonspecific variation in RNA-seq data, miCLIP provides more precise positioning by detecting exact sites of antibody-induced crosslinks (see **Fig. 2b, e** and **Extended Data Fig. 5c**, **6b**).

We additionally re-examined the Li et al. high-resolution m^1^A mapping dataset in HEK293T cells^16^. This study mapped 474 m^1^A sites in nuclear-encoded genes based on m^1^A-induced reverse transcriptase misincorporations^16^. However, upon careful examination, we eliminated 122 of these sites for the following reasons: Three sites had gene identifiers missing or removed from Refseq, 37 sites did not map to adenosines and 82 sites were duplicate sites resulting from mapping to transcript isoforms of the same gene. This left 352 unique putative m^1^A sites in nuclear-encoded genes (see **Methods** and **Extended Data Fig. 6c**).

The Li *et al.* study found most of their identified m^1^A sites occurred in the 5’ UTR. Only 19 sites were annotated at position 1 of the transcript (i.e. the transcriptional start site, TSS). However, mRNAs can have alternative transcription start sites other than (or in addition to) the RefSeq annotated transcription start site^29^. To determine the extent to which the Li *et al.* m^1^A sites mapped to alternative transcription start sites, we compared them to a list of experimentally validated transcription start sites in HEK 293T cells. This list was derived from CAGE-seq and m^6^A_m_ mapping data^30,31^. We found 134 of the putative 352 m^1^A sites overlapped with CAGE and/or m^6^A_m_ inferred transcription start sites (**Extended Data Fig. 6c**). Hence, including the 19 sites originally annotated in the Li *et al.* study, 140 m^1^A sites (or ~40%) occurred at transcription start sites. The false positive rates of these m^1^A mapping methods are not known, so it is possible that other 5’ UTR m^1^A sites are either false positives or map to currently unannotated transcription start sites. In all, our data suggest that most, if not all, of the putative start codon/5’ UTR m^1^A sites mapped by Li *et al.* are simply localized to alternative transcription start sites, consistent with our mapping results and consistent with the cap-binding properties of the m^1^A antibody.

## Discussion

Considerable attention has revolved around m^1^A as a highly prevalent mRNA modification based on its description as a high-stoichiometry, translation-promoting modification in thousands of mRNAs around start codons^3^. Subsequent studies concluded that m^1^A is less prevalent (~700 sites) with a small fraction in mitochondrial mRNA^16^, while other studies suggest that only two mRNAs have an m^1^A at a stoichiometry above 5%^5^. To address these discrepancies, we developed two m^1^A mapping approaches: (1) misincorporation mapping, a computational approach to discover m^1^A-induced misincorporations in ultra-deep RNA-Seq datasets; and (2) m^1^A miCLIP, a high-resolution method for mapping m^1^A antibody-binding sites in the transcriptome using two different m^1^A-binding antibodies. Misincorporation mapping shows that m^1^A is present at detectable stoichiometries only in the *MT-ND5* transcript, with no m^1^A in other mitochondrial mRNAs or 5’ UTRs of mRNAs as reported previously. Using m^1^A miCLIP, we find that the previously observed binding of the MBL m^1^A antibody to transcription start nucleotides and the vicinity of start codons is due to a previously unrecognized cross-reactivity of the MBL m^1^A antibody to cap structures. We confirm this using a separate m^1^A antibody (Abcam) that lacks this cap-binding cross-reactivity. We further show that m^1^A is not detectable at transcription start nucleotides, as previously proposed^16^, based on mass spectrometry and TLC analysis. Overall, these data show that the divergent m^1^A mapping data and large number of 5’ UTR-mapped m^1^A sites largely reflect cross-reactivity of the m^1^A antibody with mRNA caps.

Although the structural basis for the binding of the MBL m^1^A antibody to both m^1^A and the extended cap structure m^7^G-ppp-A will require structural studies, it is noteworthy that both these structures contain a positively charged purine; i.e., m^1^A and m^7^G. This common structural feature may be recognized by the antibody. Importantly, since the Abcam m^1^A antibody binds m^1^A but not cap structures, this antibody allows m^1^A sites to be detected without unwanted cross-reactivity to the cap. m^1^A miCLIP using the Abcam antibody shows an m^1^A signature in *MT-ND5* but does not show a transcriptome-wide enrichment in 5’ UTRs. This data, along with the mass spectrometry, TLC, and misincorporation mapping data, support the idea that the m^1^A localization to 5’ UTR sites is an artifact of the antibody rather than a reflection of *bona fide* m^1^A nucleotides at these sites.

Our results highlight how the library preparation strategies can cause an antibody-binding site at the transcription start to appear in the 5’ UTR and near start codons. The first two m^1^A mapping studies by Dominissini *et al.* and Li *et al.* mapped m^1^A to start codons and the 5’ UTR^3,4^. In mostly the same mRNAs, our miCLIP-based study mapped antibody-binding sites to transcription start nucleotides. The different location of the mapped sites reflect the different library preparation strategies used in these methods. The miCLIP strategy preserves the exact site of reverse transcription termination, which revealed binding of the m^1^A antibody to extended cap structures. In contrast, the Dominissini *et al.* and Li *et al.* m^1^A mapping studies used a library strategy that did not preserve the termination sites. Instead, the library was generated in a way that causes 3’ displacement of the peaks (see **Extended Data Fig. 1a**). Thus, the start codon-proximal m^1^A sites mapped in previous studies likely reflect displaced peaks that derive from antibody binding to transcription start nucleotides.

The finding that modification-specific antibodies have additional, unanticipated specificities is not surprising based on previous characterization of an m^6^A-binding antibody^32^. This m^6^A antibody maps both m^6^A and an m^6^A-independent polypurine sequence. Similar to this m^6^A-binding antibody, the transcriptome-wide binding patterns of the MBL m^1^A antibody reveals both m^1^A and an unintended but specific m^1^A-independent interaction with extended cap structures. Therefore, any mapped antibody-binding site in the transcriptome should be directly tested in binding assays to confirm that the antibody binds in a modification-dependent manner.

Our study supports the idea that m^1^A is a rare and low stoichiometry modification except in the case of *MT-ND5* mRNA, as seen in another study^5^. Except for *MT-ND5*, we were not able to detect m^1^A in any cytosolic mRNAs or the mitochondrial mRNAs that were reported recently to contain m^1^A^16^. This likely reflects the inability of our method to detect very low stoichiometry m^1^A modifications. Other than *MT-ND5*, other mitochondrial mRNAs may have m^1^A due to spurious low-level methylation due to their colocalization with tRNA-modifying enzymes in the mitochondrial matrix.

The Dominissini *et al.* and Li *et al.* m^1^A mapping studies found that m^1^A is highly prevalent based on mass spectrometry analysis of mRNA^3,4^. However, more recent mass spectrometry experiments showed that poly(A) preparations are contaminated with tRNA and rRNA, and when these contaminants are removed, the m^1^A signal in poly(A) preparations is absent^33^. Thus, the newer mass spectrometry data is consistent with our misincorporation and miCLIP mapping studies that indicate that m^1^A is rare.

## Acknowledgements

We thank S. Schwartz for helpful discussions, and members of the Jaffrey laboratory for helpful comments and suggestions. We thank J. Mauer and S. Zaccara for experimental contributions that were not included in this manuscript. This work was supported by NIH grants R01DA037755 (S.R.J.), T32 HD060600 and UL1 TR000457 (A.G.), T32 CA062948/KL2-TR-002385 (A.O-G.), by a Postdoctoral Enrichment Program Award from the Burroughs Wellcome Fund (A.O-G.).

## Author Contributions

S.R.J., A.V.G., and A.O.-G. conceived the project and designed and carried out experiments. A.O.-G. designed and implemented the misincorporation mapping method. A.V.G., and partly A.O.-G, performed m^1^A miCLIP, characterized the specificity of the antibodies, and analyzed the miCLIP data. M.S. performed mass spectrometry analysis of mRNA extended caps. J.M. and D.P.P assisted with computational analysis. X.L. performed chemical synthesis. S.R.J., A.V.G. and A.O.-G wrote the manuscript with input from all co-authors.

## Methods

### Cell lines and animals

For misincorporation mapping, an ultra-deep RNA-seq dataset that profiled RNA expression in blood mononucleocytes was used^10^. For m^1^A miCLIP, HEK 293T cells (passage 5-10, ATCC CRL-3216) or whole mouse brain (16 week age, pooled male and female brain, C57BL/6) was used. HEK 293T cells were purchased directly from ATCC but not further validated for identity or tested for mycoplasma contamination. Experiments involving the use of animals were approved by the Institutional Animal Care and Use Committee at Weill Cornell Medicine.

### Antibodies

The MBL m^1^A antibody in this study was a mouse monoclonal antibody that is intended to react with both m^1^A and the *N*^1^-methylated adenine base, based on the manufacturer’s specifications (MBL D345-3). The specificity was validated previously^3,4^ and here. The Abcam m^1^A antibody was a rabbit monoclonal generated against m^1^A (ab208196). Its specificity for m^1^A was validated in this study.

### Code availability

Custom scripts and computational code used in this study are available upon request.

### Alignment of reads for misincorporation mapping

Raw reads from the ultra-deep RNA-seq dataset used for this study^10^ were downloaded from GEO (accession code: GSE33029). This RNA-seq dataset was prepared using standard reverse transcription with SuperScript III, an enzyme expected to produce misincorporations at m^1^A positions^8^. Variants identified in the genomic DNA corresponding to this dataset were acquired from http://snyderome.stanford.edu. Coordinates of other SNPs that may be present in the DNA sequence were downloaded from the SNP database dbSNP (February 2017 build; https://www.ncbi.nlm.nih.gov/projects/SNP/). Read alignment of forward and reverse read mates was performed using STAR (version 2.5.3a) and the hg19 genome build. Alignment incorporated removal of PCR duplicates, and clipping of 10 bases on either end of each read, since the ends of Illumina reads are prone to sequencing error^34^. Only reads that mapped to a single location in the genome were used for downstream analysis. A maximum of one mismatch per read was permitted for alignment.

### Misincorporation mapping

To identify misincorporations, aligned reads were analyzed using Rsamtools Pileup (version 1.27.16). This program was used to determine the frequency of each of the four nucleotides present in mapped reads at every genomic position with read coverage. We limited our analysis to nucleotide positions with a minimum combined read depth of 500 unique reads across the 20 biological replicates to maximize sensitivity of detecting modified nucleotides. To prevent calling genomic variants and SNPs as modification-induced misincorporations, we did not analyze nucleotide positions containing variants discovered in the genomic DNA corresponding to the RNA-seq dataset, or SNPs annotated in dbSNP. Importantly, our analysis could only be performed on transcripts longer than the library insert size of ~250 bases^10^. For this reason, analysis of cytosolic tRNAs, which are ~75 nt-long RNAs that contain known conserved m^1^A residues, could not be performed. However, short RNAs generated from polycistronic transcripts, like mitochondrial tRNAs^35^, were represented in the analyzed library. To identify sites of modification throughout the transcriptome, we initially filtered for all nucleotide positions that were covered by at least 500 mapped reads and contained a 1% misincorporation rate, and that were present in at least half of biological replicates (**Extended Data Table 1**). To further obtain a high-confidence list of modification positions, we required that within the misincorporation profile at each initially identified position, a minimum of 5% of misincorporations were heterogeneous (i.e. transitions of the reference nucleotide to all three possible alternative nucleotides) in order to minimize detection of adenosine-to-inosine editing, and heterozygous alleles not reported as variants or SNPs. We chose this filter because hard stop nucleotides have been shown to cause heterogeneous misincorporations, even when one type of misincorporation is predominant^8^. This resulted in a high-confidence list of sites that were detected at known and novel modification positions (**Extended Data Table 2**).

### m^1^A miCLIP

m^1^A miCLIP was performed exactly as previously described^19,36^, with the following modifications: Total RNA from HEK 293T cells (*N* = 2 biological replicates) or whole mouse brain (*N* = 6 biological replicates) was extracted using TRIzol (ThermoFisher) and treated with RNase-free DNase I (Promega). Poly(A) RNA was isolated using one round of selection with oligo(dT)_25_ magnetic beads (New England Biolabs). This resulted in approximately 10 *μ* g of poly(A) RNA for each replicate used in this study. Poly(A) RNA was subjected to fragmentation using RNA Fragmentation Reagents (ThermoFisher) for exactly 12 min at 75°C. This fragmentation protocol is identical to the m^1^A-seq protocol previously used to map m^1^A, and has been reported not to facilitate substantial m^1^A to m^6^A rearrangement^3^. Fragmented RNA was then incubated with 10 to 15 *μ* g of m^1^A antibody per replicate and the antibody-RNA complexes were processed for crosslinking, immunoprecipitation, RNA 3’ linker ligation, purification, and reverse transcription exactly as previously described^19,36^. Following reverse transcription of purified peptide-RNA complexes, first-strand cDNA was circularized using CircLigase II ssDNA Ligase (EpiBio) to preserve the 3’ end of the cDNA, and thus, sites of m^1^A-induced terminations of reverse transcription. To generate priming sites for library amplification, the cDNA was cut in the middle of the cDNA primer sequence using a single-stranded DNA oligo complementary to this sequence and FastDigest *BamH*I (ThermoFisher) as previously described^19,36^. This generated priming sites for the Illumina P5 and P3 primers on either side of the first-strand cDNA, eliminating the need for a second-strand synthesis step. For library amplification, Accuprime Supermix I (ThermoFisher) and Illumina P5 and P3 primers were used (see **Extended Data Table 7**). PCR amplification cycle number was selected as previously described^36^. Amplified libraries were purified using AMPure XP magnetic beads (Beckman Coulter). Libraries were subjected to next-generation sequencing at the Epigenomics Core of Weill Cornell Medicine. Libraries were sequenced on an Illumina HiSeq 2500 and MiSeq instrument in single-end mode to generate 50-base reads.

### RNA-seq

HEK 293T cell total RNA was extracted with TRIzol (ThermoFisher), treated with RNase-free DNase I (Promega), and poly(A) RNA was isolated using one round of selection with oligo(dT)_25_ magnetic beads (New England Biolabs). RNA was then subjected to fragmentation using RNA Fragmentation Reagents (ThermoFisher) for exactly 12 min at 75°C. Fragmented RNA was then subjected to RNA 3’ linker ligation using T4 RNA Ligase I (New England Biolabs) and reverse transcription using a primer complementary to the linker sequence and SuperScript III (ThermoFisher) (see **Extended Data Table 7**). First-strand cDNA was gel-purified using denaturing PAGE, and then circularized using Circligase II ssDNA Ligase (EpiBio). Circularized cDNA was then cut and amplified exactly as described above for m^1^A miCLIP. Resulting RNA-seq libraries were subjected to next-generation sequencing at the Epigenomics Core of Weill Cornell Medicine. Libraries were sequenced on an Illumina HiSeq 2500 instrument in single-end mode to generate 50-base reads.

### Read processing and alignment

After sequencing, reads from m^1^A miCLIP or RNA-seq libraries were trimmed of the 3’ linker sequence and barcoded reverse transcription primer sequences using Flexbar (version 2.5) (see **Extended Data Table 7**). To demultiplex reads belonging to individual biological replicates, the pyBarcodeFilter.py script of the pyCRAC suite (version 1.2.2) was used. The random portion of the reverse transcription barcode was then moved into the sequence header using a custom awk script (available upon request). PCR duplicates were collapsed using pyFastqDuplicateRemover.py of the pyCRAC suite. Finally, reads were aligned to hg19 for HEK 293T cells or mm10 for mouse brain using Bowtie (version 1.1.2).

### Generation of m^1^A miCLIP clusters

m^1^A miCLIP clusters of unique reads were generated using the CIMS software package for analysis of HITS-CLIP data^37,38^. To generate clusters and determine the cluster score (maximum of stacked reads), the tag2profile.pl, tag2cluster.pl, and extractPeak.pl scripts of the CIMS software package were used. A custom awk script was then used to filter for clusters of a minimum score (at least 20 stacked reads; script is available upon request).

### Motif analyses

To search for a possible common sequence motif present in our HEK 293T cell m^1^A miCLIP dataset, we focused on potential motifs present in m^1^A miCLIP clusters in the 5’ UTR, the region of predominant m^1^A miCLIP cluster enrichment. The genomic sequences of these clusters were retrieved using bedtools and subjected to motif discovery using the MEME suite (version 4.11.4).

### Metagene distribution analyses

To analyze the metagene distribution of m^1^A miCLIP clusters on mRNAs, MetaPlotR was used^39^, with in-house modifications. The density of m^1^A miCLIP coverage was normalized to that of RNA-seq coverage to reveal any enrichments using a custom R script (available upon request). For the HEK 293T cell metagene, the in-house HEK 293T cell RNA-seq dataset described above was used for normalization. For the mouse brain metagene, a published whole-brain RNA-seq dataset was used^40^ (accession code: GSE52564). To plot the coverage of transcription start sites by m^1^A miCLIP at higher resolution, the plotProfile tool of the Deeptools suite was used.

### Examination of antibody crosslink sites around the transcription start site

For analysis of antibody crosslinks at the transcription start sites of mRNAs, we analyzed terminations of reverse transcription (i.e. 5’ ends of reads) around these sites. To do so, the number of terminations was measured around RefSeq-annotated transcription start sites that had coverage in both m^1^A miCLIP and RNA-seq. Terminations were counted at positions ranging from the transcription start site to position +4 relative to the transcription start site. Then, transcription start sites were filtered for those that contained a minimum coverage of 5 unique reads at the transcription start site position in both m^1^A miCLIP and RNA-seq. This filtered set of transcription start sites was then used to compare the distributions of read terminations in m^1^A miCLIP and RNA-seq. We focused on terminations rather than misincorporations in m^1^A miCLIP is because the misincorporation profile of m^1^A is sequence-dependent, with both upstream and downstream nucleotides contributing to misincorporation variability^8^. Thus, we used the presence of terminations as a signature of antibody crosslinking events in our dataset. Additionally, while rare types of reverse transcriptases that read through m^1^A have been described^41^, standard reverse transcriptases, like the SuperScript III used in m^1^A miCLIP, produce frequent terminations at m^1^A residues^8,42^.

### Measurement of transcription start sites enriched by m^1^A miCLIP

To determine the types of transcription start sites overlapping m^1^A miCLIP clusters, we used a collection of transcription start sites that included RefSeq transcription start sites as well as recently-mapped transcription start site regions containing the m^6^Am mRNA extended cap^23^. The frequencies of all transcription start site types or those overlapping m^1^A miCLIP clusters were thus determined using this collective set.

### Synthetic oligonucleotides used in this study

For biochemical analysis of various modifications present within the extended caps of mRNAs, synthetic oligonucleotides were generated as standards for mass spectrometry and/or thin-layer chromatography (see below; see **Extended Data Table 7**). Oligonucleotides containing m^7^G-ppp-A_m_, m^7^G-ppp-m^6^A, or m^7^G-ppp-m^6^A_m_ were synthesized chemically as described previously^43^. Oligonucleotides containing m^7^G-ppp-m^1^A or m^7^G-ppp-m^1^A_m_ were synthesized enzymatically using an oligonucleotide initiating with ppp-m^1^A (Trilink). This oligonucleotide was capped using ScriptCap Cap 1 Capping System (CellScript) to generate the m^7^G cap and, in the case of m^7^G-ppp-m^1^A_m_, 2’-*O*-methylation of m^1^A.

### Liquid chromatography and mass spectrometry (LC-MS)

Poly(A) RNA was prepared for mass spectrometry as follows. Total RNA from HEK 293T cells was treated with TURBO DNase (ThermoFisher) according to the manufacturer’s instructions, followed by two rounds of poly(A) selection using oligo(dT) magnetic beads (NEB). Small RNAs shorter than 200 nt were then removed from the poly(A) RNA using the RNeasy kit (Qiagen). This size selection was performed to prevent detection of extended cap structures that are known to be present in certain small RNAs, like small nuclear RNAs (snRNAs). Approximately 5 *μ* g of DNase-treated, poly(A)-selected, and size-selected RNA was thus generated for each sample for mass spectrometry analysis. To release extended cap structures from the nucleotides comprising the internal portion of the RNA, RNA was digested with 2-4 units of Nuclease P1 (Sigma Aldrich) in a final buffer concentration of 30 mM sodium acetate (pH 5.5) for 3 hours at 37°C. Following digestion, the nuclease was removed from the samples using molecular weight cutoff centrifugal filters (VWR). The digested and purified RNA was finally dried using an Eppendorf Vacufuge and reconstituted with 70% acetonitrile (LC-MS grade; Sigma Aldrich) to a final concentration of 0.5 μg/μl. 2 μl of the resulting solution were subjected to MS analysis.

Samples were injected into an LC-MS/MS system comprised of an Agilent 1260 HPLC and an Agilent 6460 triple quadrupole mass spectrometer equipped with a JetStream electrospray ionization source. Positive ion monitoring and multiple reaction monitoring was used for detection of extended caps. The caps were resolved on an aqueous normal phase column (ANP, Cogent Diamond Hydride, 4 μm particle size, 150 mm × 2.1 mm; Microsolv). To achieve chromatographic separation of the cap structures from mononucleotides, the following gradient was used. The aqueous mobile phase (Buffer A) was 50% isopropanol with 0.025% acetic acid, and the organic mobile phase (Buffer B) was 90% acetonitrile containing 5 mM ammonium acetate. EDTA was added to the mobile phase in a final concentration of 6 μM. The final gradient applied was: 0-1.0 min 99% B, 1.0-7.0 min to 80% B, 7.0-18.0 min to 50% B, 18.0-19.0 min to 0% B and 19.1 to 29.0 min 99% B. The flow rate was 0.4 mL/min during data acquisition and 0.6 mL/min during column re-equilibration. Data was saved in centroid mode using MassHunter workstation acquisition software (Agilent). Data files were processed with MassHunter Qualitative Analysis Software (Agilent).

Exact operating source parameters for the LC-MS analysis are available upon request.

### Biochemical examination of modifications at transcription start sites

To identify the initiating nucleotide structure in mRNAs bound by the m^1^A antibody, we utilized two dimensional thin layer chromatography (2D-TLC) analysis of mRNA extended caps as previously described^23^. For analysis of cellular RNA, we used HEK 293T cell poly(A) RNA or poly(A) RNA enriched using the m^1^A antibody as previously described^3,4^. Oligonucleotide standards, input (antibody-unbound) poly(A) RNA, and m^1^A antibody-enriched poly(A) RNA were then subjected to 2D-TLC as previously described^23^, with one modification. To enhance resolution of m^1^A and m^1^A_m_ from other nucleotide species, the second dimension of 2D-TLC resolution was performed using the following buffer: 60% ammonium sulfate in 100 mM sodium phosphate buffer of pH 6.8 (w/v), and a final concentration of 2% *n*-propanol.

### Synthesis of *N*^1,6^-methyladenosine

To synthesize *N*^1,6^-methyladenosine, *N*^6^-methyladenosine (Selleckchem) was dissolved in dry DMF and followed with addition of iodomethane (Acros Organics; 10:1 molar ratio iodomethane:*N*^6^-methyladenosine). The mixture was stirred overnight at room temperature. The product was purified by flash chromatography on silica gel (EMD), eluting with methanol and dichloromethane (1:10 to 1:5; ACS or HPLC grade solvents). This resulted in a product yield of 46.3% *N*^1,6^-methyladenosine. Product identity was confirmed by nuclear magnetic resonance (NMR) and high resolution mass spectrometry (HR-MS).

NMR spectra were recorded using a 500-MHz Bruker DMX-500 instrument at room temperature, and chemical shifts were referenced to the residual solvent peak. Shifts were as follows: ^1^H NMR (500 MHz, DMSO-*d*_6_) δ 8.11 (s, 1H), 8.04 (s, 1H), 5.76 (d, *J* = 5.8 Hz, 1H), 5.15 (s, 1H), 4.43 (t, *J* = 5.4 Hz, 1H), 4.15 (m, 2H), 3.92 (d, *J* = 3.7 Hz, 1H), 3.63 (dd, *J* = 12.0, 3.9 Hz, 1H), 3.56 (m, 2H), 3.50 (s, 3H), 3.45 (d, *J* = 6.5 Hz, 1H), 1.23 (s, 3H).

HR-MS data was recorded with Waters LCT-Premier XE at room temperature. For a predicted mass for *N*^1,6^-methyladenosine, or C_12_H_18_N_5_O_4_^+^, of 296.1353, the mass found was 296.1361.

### Analysis of Li *et al.* m^1^A sites

The Li *et al.* m^1^A sites for nuclear-encoded genes was obtained from supplemental Table S2 of the published manuscript. We annotated each of the 474 transcriptomic sites with their corresponding genomic coordinates and nucleotide sequences using an annotation file generated from Refseq with MetaPlotR. With a custom R script, we then filtered likely erroneous sites as specified in the Results section. Briefly, sites corresponding to gene IDs missing in Refseq, or that mapped to non-adenosine nucleotides, or with duplicate genomic coordinates, were all removed.

### Characterization of antibody affinity for various substrates

The specificity or affinity of the m^1^A antibody for various nucleosides, nucleotides, or cap structures was determined as follows. To determine the specificity of the m^1^A antibody for various nucleosides, two approaches were performed. For testing the specificity of the antibody for various nucleosides in the context of m^1^A miCLIP, the antibody was crosslinked to total cellular RNA in the presence of various competitor nucleotides. Antibody binding, crosslinking, and detection of crosslinked antibody-RNA complexes was performed exactly as in miCLIP, except with the inclusion of the competitor nucleotide during the antibody binding reaction.

For testing the specificity of the antibody for modified adenines, especially those resembling *N*^1^-methylated adenine, a previously described dot blot assay was performed wherein the competing molecule is added during antibody binding^1^. Competition assays are usually used to measure binding, rather than spotting the nucleotides to the membrane, since the manner of interaction of each nucleotide to the membrane is not known and can affect antibody binding. For measuring the affinity, the IC_50_ of the various nucleotides and cap dinucleotides was measured. In these experiments, a series of the dot blot assays were performed, where serial dilutions of each competitor molecule (ranging from 10 μM to 1 nM) were used in parallel during antibody binding reactions. The dot blots were performed as follows: 250 ng of m^1^A-containing synthetic oligonucleotide (see **Extended Data Table 7**) were spotted in triplicate on a BrightStar membrane (ThermoFisher), allowed to briefly air-dry, and auto-crosslinked twice in a Stratalinker 2400 (Stratagene). Each membrane was rinsed briefly in PBST, then blocked for 1 h at room temperature in 5% milk in PBST. Each membrane was then placed into a pouch containing a 1:1000 dilution of the m^1^A antibody in 0.5% milk in PBST, and an appropriate concentration of competitor molecule. The antibody binding proceeded for 2 h at room temperature. Then, each membrane was washed three times in PBST (5 min per wash), and then incubated in a dilution of 1:2500 of secondary antibody (anti-mouse, GE # NA931; anti-rabbit, GE NA934) in 0.5% milk in PBST for 1 h at room temperature. Finally, the membrane was washed three times in PBST (5 min per wash), and developed using ECL Prime (GE). Membranes corresponding to a dilution series of a specific competitor molecule were imaged together using a ChemiDoc Imager (Bio-Rad).

## Extended Data Figure Legends

**Extended Data Figure 1.**
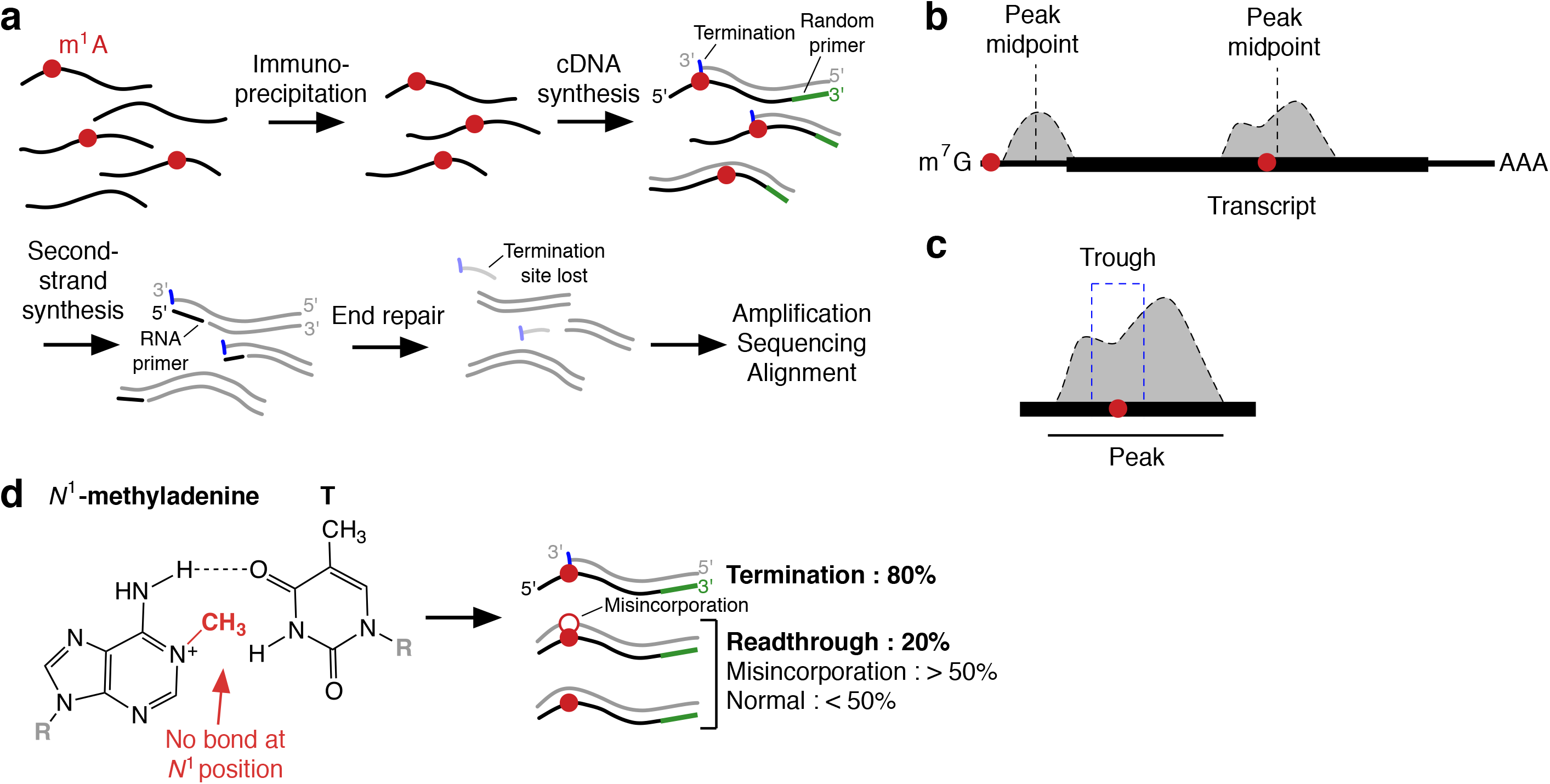
m^1^A mapping by immunoprecipitation and its effect on base-pairing. **a**, Earlier strategy for mapping m^1^A. Earlier transcriptome-wide m^1^A mapping approaches^3,4^ generated sequencing libraries from RNA fragments containing m^1^A. To achieve this, RNA (black lines) was fragmented, immunoprecipitated using a m^1^A antibody, and reverse transcribed using primers (green bars). m^1^A (red circles) predominantly causes termination of standard reverse transcription, and only a small fraction of cDNAs (grey lines) go through the m^1^A residue. Although the termination site marks the position of the m^1^A, the exact site of the reverse transcription termination is not preserved in these methods. This is because during cloning of the cDNA library, the second-strand cDNA is generated in a manner that uses RNase-H to nick the RNA template. This RNAse-H-generated nick produces an RNA primer (short black line) for second-strand synthesis. Following second-strand synthesis, the RNA primer and the 3’ end of the first-strand cDNA are lost during end repair (i.e. blunting) of double-stranded cDNA ends. Thus, in earlier m^1^A mapping approaches, the site where the reverse transcriptase terminates is lost, preventing m^1^A residues from being detected at nucleotide resolution. **b**, Peaks generated by earlier m^1^A mapping approaches can be displaced from the m^1^A site. Although the precise sites of reverse transcription terminations are not preserved in earlier m^1^A mapping approaches, peaks (grey) were used to infer the presence of m^1^A (red circles) in transcripts (black bar). If the m^1^A is at an internal site, reads will be generated on either side of the m^1^A. The resulting peak is likely to cover the m^1^A site (right). The “peak midpoint,” which was previously reported for these peaks^3^, is likely to be near the m^1^A site. A different result happens if the modification is at the 5’ end of mRNAs (left). In this case, because cDNA ends are not preserved (see **a**), peaks will be displaced downstream of the m^1^A site and the peak midpoint will not reflect the location of the modified residue. Since the cDNA termination site is lost, there will be no way to identify modified sites at the 5’ ends of transcripts. In contrast, m^1^A miCLIP can detect either an both internal and 5’ terminal modifications of the mRNA since the termination sites are preserved. **c**, Peak troughs proposed to enhance m^1^A mapping resolution in earlier approaches^3^. If an m^1^A is located at an internal site in the mRNA (i.e. not at the transcription start site), regions containing m^1^A residues (red circle) can be narrowed down based on the regions of reduced read coverage, or “troughs,” (box) within peaks (grey). Troughs are thought to occur at m^1^A residues because cDNA generated by reverse transcription can be primed upstream of the m^1^A residue, or downstream of the m^1^A residue. However, troughs are a relatively nonspecific feature of RNA-seq and are especially difficult to distinguish from noise in low abundance or low coverage mRNAs. **d**, Mechanism of the effect of *N*^1^-methylated adenine on reverse transcription. *N*^1^-methylated adenine leads to termination of reverse transcription or misincorporation of nucleotides because the *N*^1^ methyl on the adenine base blocks formation of a hydrogen bond that is necessary to pair with thymine. As a result, *N*^1^-methylated adenine and m^1^A cause termination of reverse transcription, and any readthrough of reverse transcriptase through this modification is often accompanied by misincorporation of the complementary nucleotide.

**Extended Data Figure 2.**
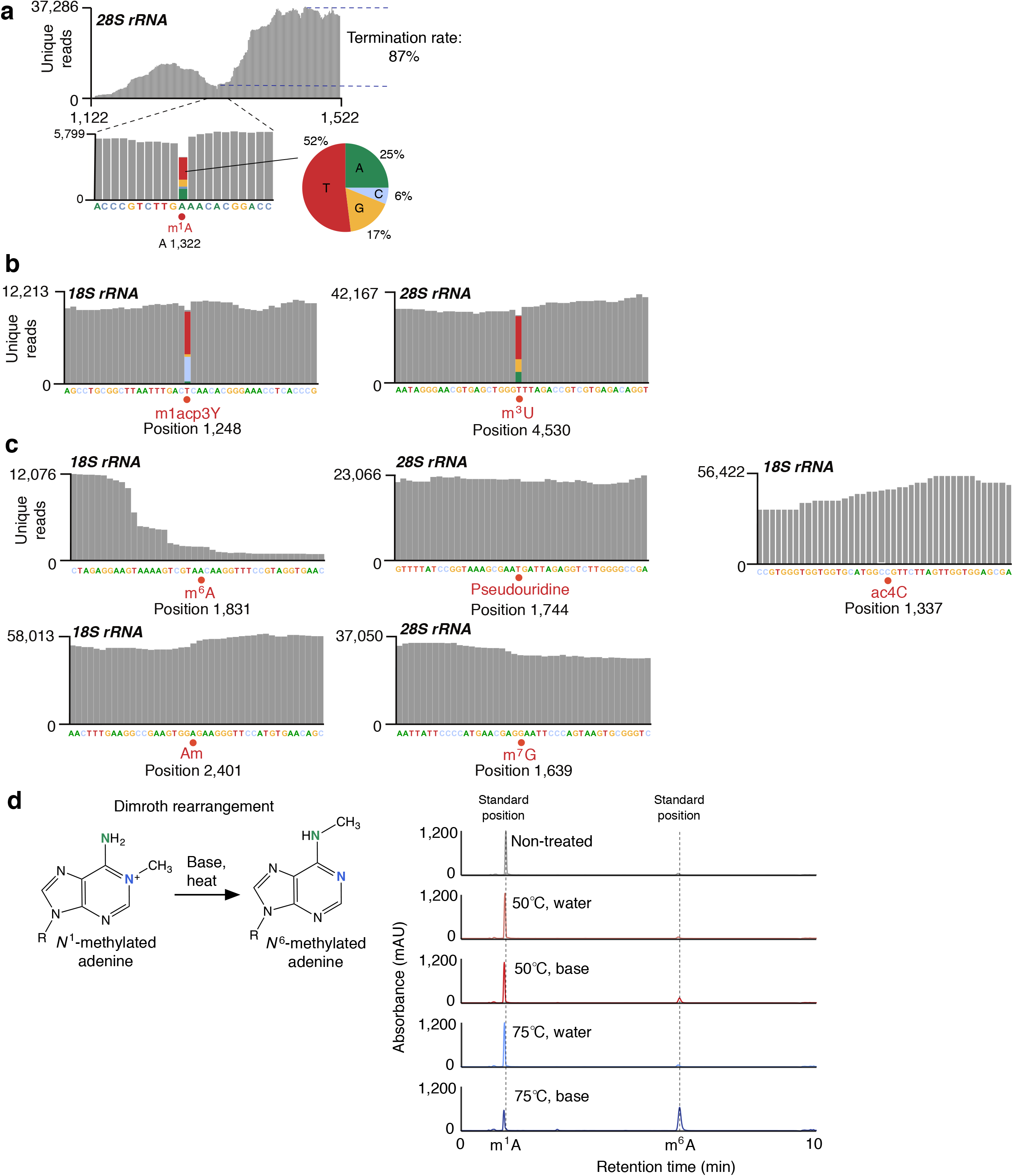
Validation of misincorporation mapping. **a**, Misincorporation mapping detects m^1^A in the *28S* rRNA. Shown is the quantification of misincorporations described in **Fig. 1b**. As expected, the m^1^A residue at this position was marked by a high rate of misincorporations. 17% of the reads (grey) at this position showed an A→G transition, 6% of reads contained an A→C transition and most reads—52%—contained an A→T transition. Together, this misincorporation profile is characteristic of the reverse transcription signature of m^1^A^8^. The detection of misincorporations was important for our study because the RNA-seq dataset we used for misincorporation mapping was cloned in a manner that did not preserve m^1^A-induced reverse transcription terminations^10^. Thus, we relied on misincorporations to map hard stop modifications like m^1^A. This result demonstrates that misincorporation mapping can be used to map m^1^A residues using a misincorporation profile that is typical of m^1^A. We considered that in our misincorporation mapping dataset, Dimroth rearrangement could have occurred, preventing the detection of m^1^A residues. However, given our ability to identify m^1^A in the *28S* rRNA using this strategy, this appears unlikely. To estimate how much m^1^A was lost during library preparation, we measured the termination rate seen in the *28S* rRNA. As can be seen, there is high coverage downstream of the m^1^A. This drops substantially at the m^1^A site, reflecting a read termination rate of ~87%. This is consistent with a near complete preservation of m^1^A in this RNA. Thus, we reasoned that Dimroth rearrangement would not substantially impair our ability to detect m^1^A in this dataset. **b**, Misincorporation mapping detects other hard stop modifications. In addition to detecting the m^1^A residue in the *28S* rRNA, misincorporation mapping reliably identified other hard stop modifications in rRNA. In the *18S* rRNA, 1-methyl-3-(3-amino-3-carboxypropyl)pseudouridine (m^1^Acp3Y) was marked by reads (grey) containing a high rate of misincorporations. Likewise, another hard stop modification, m^3^U, in the *28S* rRNA, was also marked by misincorporations. Thus, misincorporation mapping can identify a variety of hard stop modifications. **c**, Misincorporation mapping does not detect modifications that do not affect reverse transcription. To validate that modification mapping specifically detected hard stop modifications, we examined the misincorporation profiles of modifications in rRNA that are known not to affect reverse transcription. As expected, we found that four modifications that do not affect reverse transcription, including m^6^A and 2’-*O*-methyladenosine (A_m_) in the *18S* rRNA, and pseudouridine and m^7^G in the *28S* rRNA, produced no misincorporations in the reads (grey) at their respective positions. Thus, misincorporation mapping is specific to hard stop modifications, and not modifications that do not affect reverse transcription. **d**, m^1^A is relatively stable under routine laboratory conditions, but unstable in the presence of base and heating. Shown is the m^1^A to m^6^A conversion via the Dimroth rearrangement (left mechanism). Under basic conditions and high temperatures, m^1^A can convert to m^6^A (right). Shown is an HPLC assay to detect conversion of m^1^A (0.5 mM) to m^6^A in the following conditions: 25^°^C (water, 30 min), 50^°^C (water, 30 min), 75^°^C (water, 30 min), 50^°^C (10 mM sodium bicarbonate, pH 9.0, 30 min), 75^°^C (10 mM sodium bicarbonate, pH 9.0, 30 min). Minimal conversion was seen except at 75^°^C in basic conditions. Under these conditions, ~50% conversion was seen. HPLC assay conditions were as previously described^23^.

**Extended Data Figure 3.**
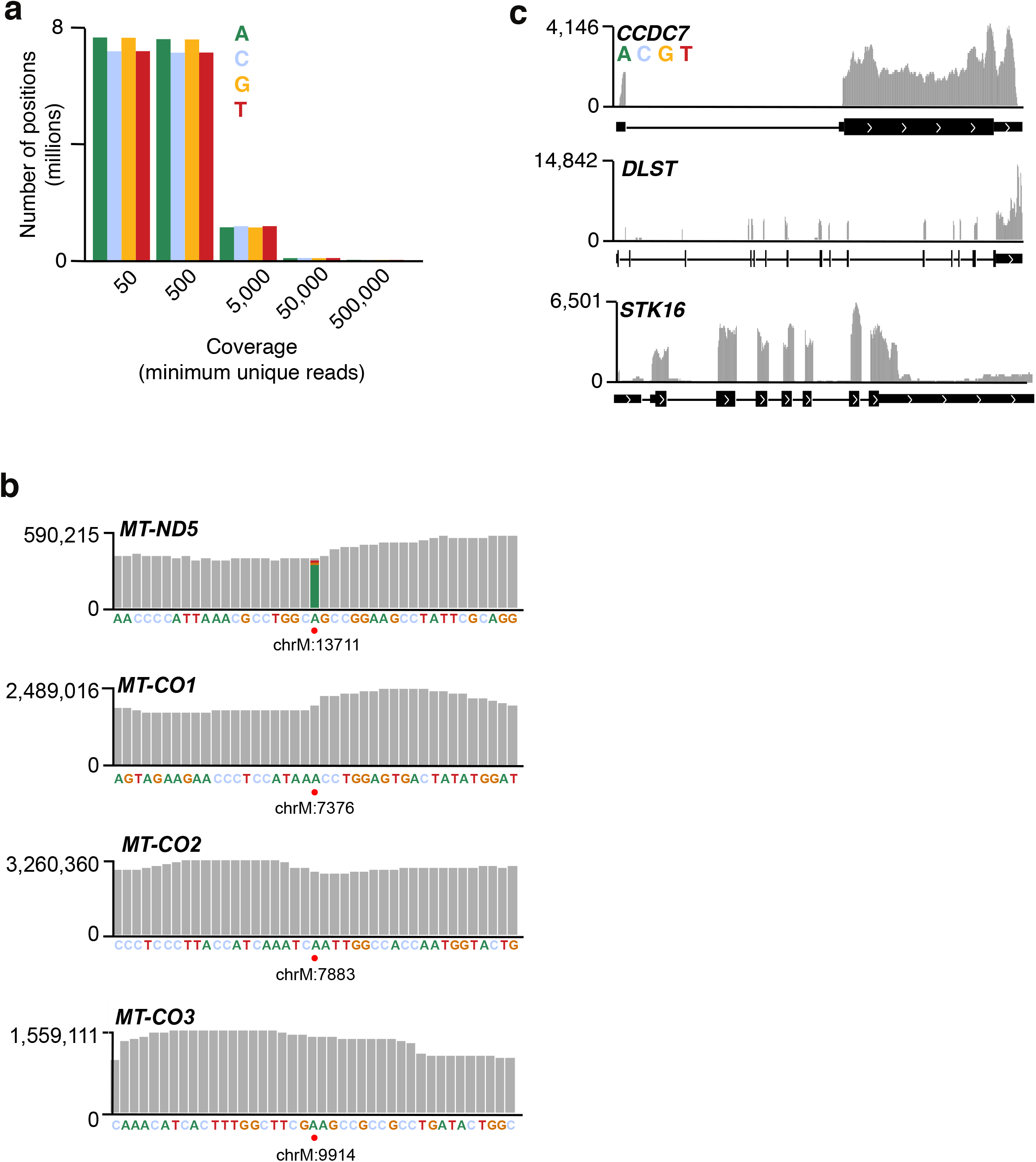
Misincorporation mapping in mRNA. **a**, Selection of read depth criteria for misincorporation mapping. To detect modification-induced misincorporations throughout the transcriptome, we first examined the coverage depth of the ultra-deep RNA-seq dataset at all genomic positions. At positions that had coverage, many positions were covered by 50 to 500 unique reads. However, for our analysis, we chose to analyze the subset of nucleotides that were covered by 500 or more unique reads in order to maximize the sensitivity of detecting hard stop modifications in mRNA. Here, in total, approximately 8 million positions of each type of nucleotide were covered by 500 or more reads. Thus, many nucleotides throughout the transcriptome were able to be evaluated for hard stop modifications. **b**, Examination of misincorporations in putative m^1^A-containing mitochondrial transcripts. The indicated positions in *MT-ND5*, *MT-CO1*, *MT-CO2* and *MT-CO3* were identified as putative m^1^A sites in two previous studies. A robust m^1^A misincorporation signature was seen for the *MT-ND5* site but not at the m^1^A sites in the other genes. Thus, with the exception of *MT-ND5*, misincorporation mapping indicates that these mRNAs lack high stoichiometry m^1^A sites. Of note, the read coverage in all genes was exceptionally high, allowing for sufficient read depth needed to identify low stoichiometry m^1^A sites. The exact misincorporation rate at each m^1^A site is listed in **Extended Data Table 3** for each putative m^1^A-containing mitochondrial mRNA. Position of putative m^1^A sites for each transcript is indicated with a red dot. **c**, Examination of misincorporations on high-stoichiometry m^1^A-containing transcripts. Previously identified m^1^A containing transcripts, *CCDC7*, *DLST*, and *STK16*, were shown in an earlier study to contain m^1^A residues of greater than 50% stoichiometry. However, no adenosines in these transcripts showed misincorporation profiles that met our filtering criteria (see **Methods**), despite high depth of coverage (black, transcript models; white arrows, direction of transcription). Thus, misincorporation mapping did not detect m^1^A residues in the transcripts previously reported to contain m^1^A at high stoichiometry.

**Extended Data Figure 4.**
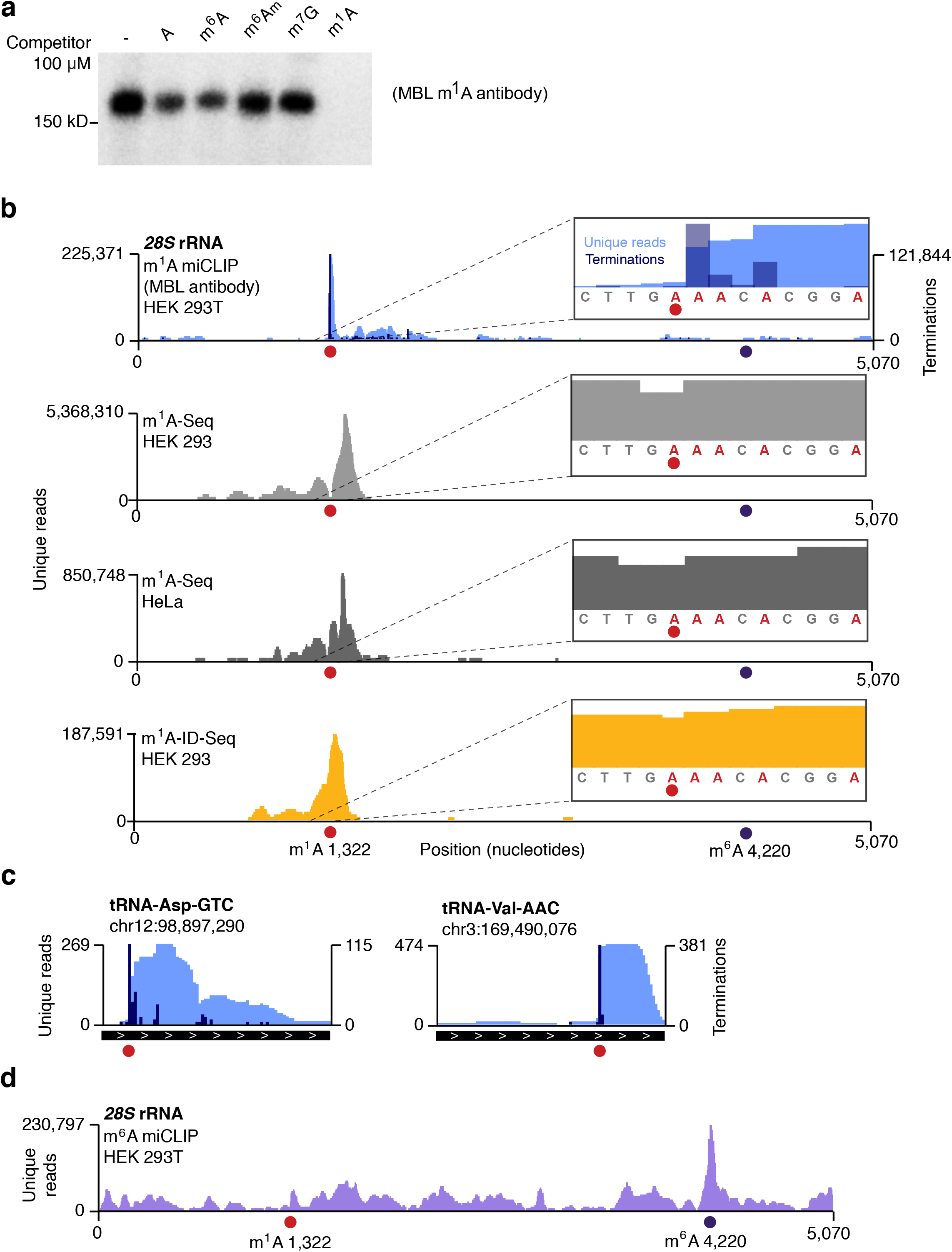
m^1^A miCLIP validation with the MBL m^1^A antibody. **a**, The MBL m^1^A antibody detects m^1^A RNA. Although the specificity of the m^1^A antibody was previously validated^3,4^, we wanted to validate its specificity in the context of the m^1^A miCLIP protocol. To do this, we incubated the antibody with total fragmented RNA from HEK 293T cells in the presence of 100 μM of various competitor nucleotides. Then, the antibody was crosslinked to the RNA and the crosslinked antibody-RNA complexes were radiolabeled and purified by SDS-PAGE and membrane transfer. While some competitor nucleotides weakly inhibited antibody-RNA crosslinking, only m^1^A abolished all crosslinking. Moreover, we did not readily detect binding of the antibody to these other nucleotides when the RNA was subjected to sequencing (see **b**). **b**, m^1^A miCLIP detects the m^1^A residue in the *28S* rRNA. To further assess the specificity of m^1^A miCLIP, we aligned unique reads from m^1^A miCLIP (light blue) or earlier m^1^A mapping approaches (m^1^A-seq^3^, grey; m^1^A-ID-seq^4^, yellow) to rRNA. Like earlier m^1^A mapping approaches, the m^1^A miCLIP protocol resulted a striking accumulation of unique reads at the m^1^A residue at position 1,322 (red circle) of the *28S* rRNA. Notably, no enrichment of reads was observed at the m^6^A residue at position 4,220 (purple circle), or anywhere else, confirming a lack of cross-reactivity of the antibody to *N*^6^-methylated adenine or other modified nucleotides, which are abundant in rRNA. Moreover, while all protocols resulted in read accumulations at the m^1^A residue, the m^1^A miCLIP peak was markedly narrower than the peaks produced by earlier strategies. Furthermore, in m^1^A miCLIP, this m^1^A residue was marked by many read terminations (dark blue) at the +1 position of the m^1^A residue. Thus, m^1^A miCLIP marks m^1^A residues with high specificity and resolution. **c**, m^1^A miCLIP detects m^1^A in tRNAs. To validate the specificity of m^1^A miCLIP further, we examined tRNAs (black bars; white arrows, direction of transcription), which contain conserved m^1^A residues (red circles, MODOMICS annotations^45^). Here, two example tRNAs are shown. Like in the *28S* rRNA, in these tRNAs, m^1^A miCLIP reads (light blue) and terminations (dark blue) marked m^1^A residues with high specificity and resolution. **d**, miCLIP does not cause rearrangement of m^1^A to m^6^A. To determine whether the miCLIP protocol may miss detection of m^1^A residues due to Dimroth rearrangement of m^1^A to m^6^A, we examined the presence of potential m^6^A at the *28S* rRNA m^1^A residue position. While the conserved m^6^A residue was marked by a high enrichment of accumulated m^6^A miCLIP reads (purple circle), reads found at the position of the m^1^A residue (red circle) were present at background level. This suggests minimal conversion of m^1^A to m^6^A during the miCLIP protocol.

**Extended Data Figure 5.**
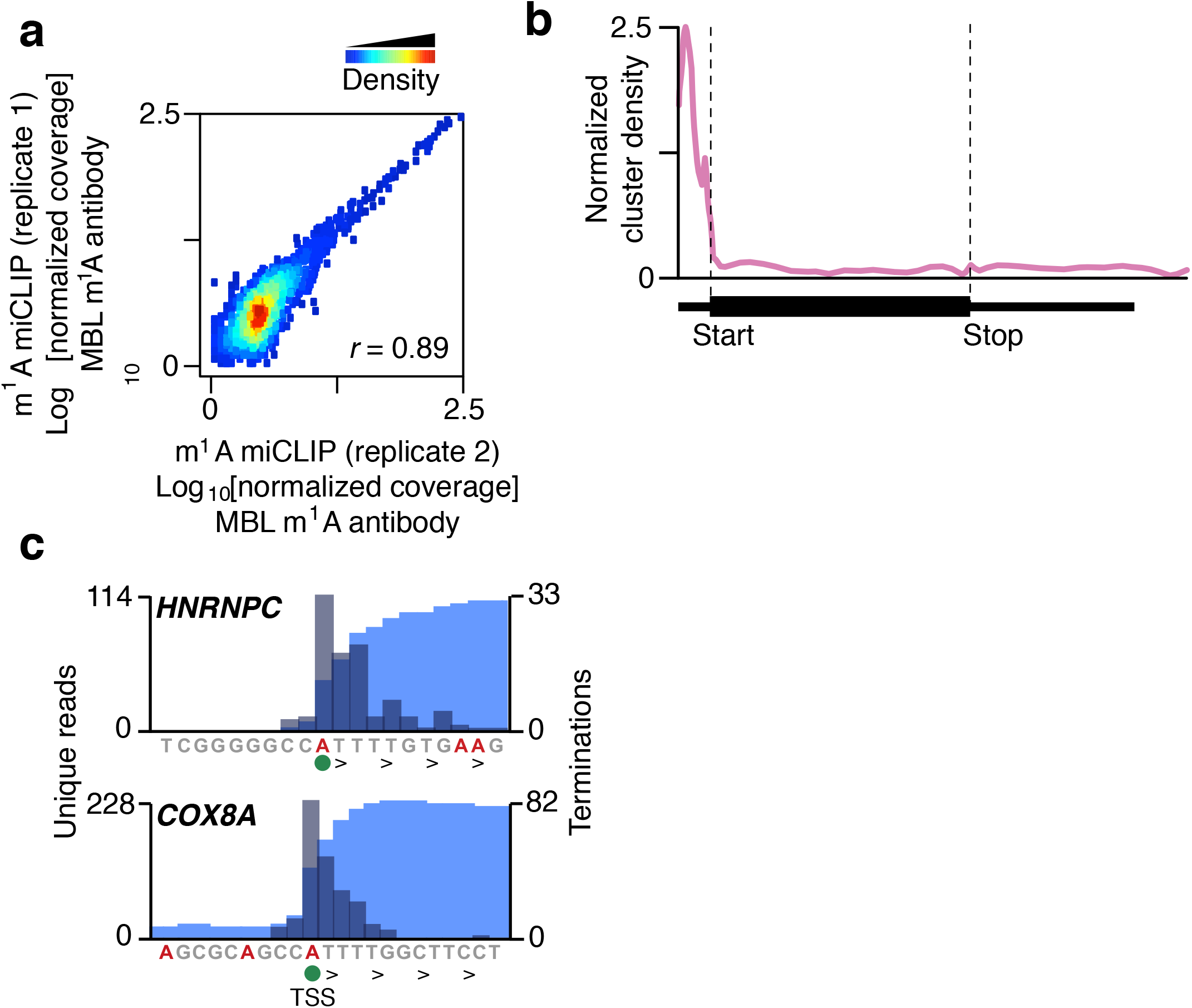
m^1^A miCLIP with the MBL antibody enriches for reads close to transcription start nucleotide. **a**, m^1^A miCLIP replicate correlation. The x and y axes of the scatter plots represent normalized read coverage in 100 nt bins on the human genome in replicate 1 (x) or replicate 2 (y) of m^1^A miCLIP performed in HEK 293T cells. The replicates show highly reproducible read coverage across the genome (*r* = 0.89, Pearson correlation). **b**, Metagene distribution of m^1^A miCLIP clusters in mouse brain transcripts. The density of m^1^A miCLIP clusters in mouse brain mRNAs was normalized to RNA-seq coverage (see **Methods**) and plotted on a virtual transcript (start, start codon; stop, stop codon). Like in human transcripts, m^1^A miCLIP clusters in mouse brain mRNA were highly enriched in the 5’ UTR, and particularly, next to the transcription start site. **c**, Characterization of m^1^A miCLIP terminations at positions +2 and +5 relative to the putative site of antibody crosslinking. While many transcripts with m^1^A miCLIP clusters (light blue) in their 5’ UTR showed enrichment of read terminations (dark blue) at the +1 position relative to the initiating nucleotide, certain transcripts had additional terminations between positions +2 and +5. We considered the possibility that these terminations reflect arrests of cDNA synthesis due to the presence of crosslinked antibody peptide on adenosine. Indeed, this was likely the case, as we observed that on certain transcripts like *HNRNPC* and *COX8A*, even though terminations occurred between positions +2 and +5 relative to the initiating adenosine, the only adenosine near terminations was at position 0 (green circles). Indeed, antibody crosslinks at transcription start sites have been previously shown to generate read terminations at positions +2 to +5 relative to the transcription start site, while the antibody crosslinking site was only at the transcription start site^19^.

**Extended Data Figure 6.**
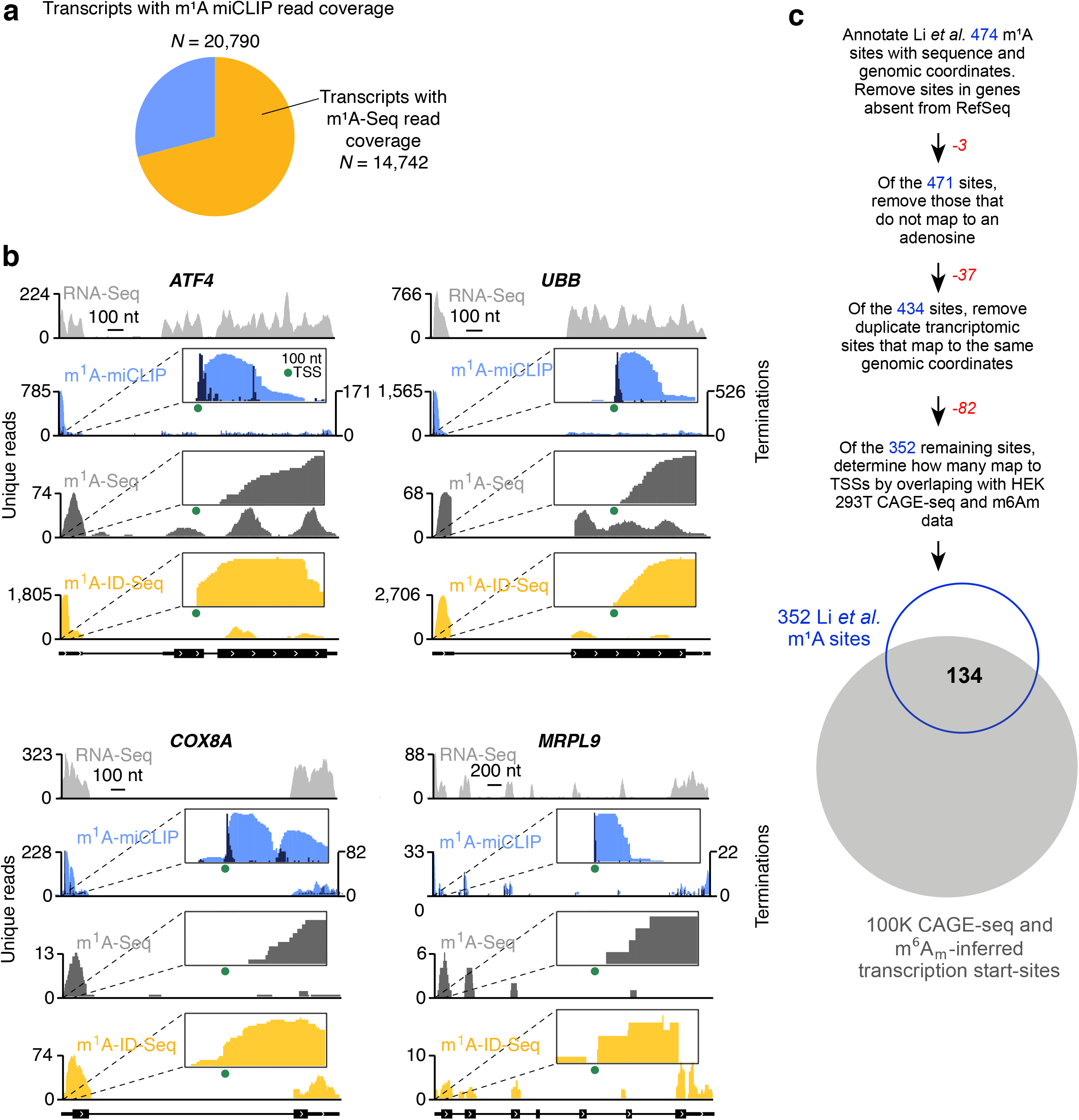
Comparison of m^1^A miCLIP (MBL antibody) and earlier m^1^A mapping strategies. **a**, m^1^A miCLIP and the earlier m^1^A-Seq mapping strategy mark similar mRNAs. To compare m^1^A miCLIP with previous maps, we first identified unique transcripts containing m^1^A miCLIP or m^1^A-Seq unique read coverage. Of the 20,790 transcripts containing m^1^A miCLIP coverage, more than ~70% also showed m^1^A-Seq coverage. This demonstrates that overall, the m^1^A miCLIP profile of the transcriptome is similar to that of m^1^A-Seq. **b**, m^1^A miCLIP read distribution differs from earlier mapping methods. Here, we wanted to demonstrate that while m^1^A miCLIP (light blue) and earlier methods (m^1^A-Seq, dark grey; m^1^A-ID-Seq, yellow) have coverage on similar transcripts, m^1^A miCLIP marks transcription start sites (green circles) due to differences in read distribution. The m^1^A miCLIP, m^1^A-Seq, and m^1^A-ID-Seq coverage of four HEK 293T transcripts, *ATF4*, *UBB*, *COX8A*, and *MRPL9* (black bars; white arrows, direction of transcription), are shown. While all four transcripts demonstrated predominant peaks in their 5’ UTRs in all three mapping protocols (see light grey RNA-Seq to note m^1^A peak enrichment), m^1^A miCLIP clearly recognized adenine at transcription start sites (see insets). This is because earlier mapping strategies did not preserve the full-length cDNA produced by reverse transcription of immunoprecipitated RNA fragments, resulting in a loss of detection of RNA 5’ ends (see **Extended Fig. 1a**). m^1^A miCLIP, on the other hand, preserves the full-length cDNA (see **Fig. 2a**), resulting in ready detection of transcription start sties. Taken together, these data demonstrate that the peaks identified by earlier m^1^A mapping approaches are likely to be derived from adenine-containing extended cap structures in mRNAs. **c**, Top, flow chart showing the filtering of non-validated m^1^A sites from the Li *et al.* study. Bottom, Venn diagram showing overlap of final list of m^1^A sites with transcriptional start sites inferred from CAGE-seq and m^6^A_m_ mapping data. Nearly 40% of putative m^1^A sites are at transcriptional start sites.

