## Supplementary material for "Antibody cross-reactivity accounts for widespread appearance of m^1^A in 5’ UTRs": code: analyze_split_rpileup_method2.pl

```
#!/usr/bin/perl
use strict;
use warnings;

my @file_list = @ARGV;
print "Number of files: ", scalar (@file_list), "\n";

my %var_h;
# Snyder called variants
# open(SVAR, "TB0001907.all.ILLUMNIA.bwa.CEU.high_coverage.20101118.snp.raw.vcf") || die "Can't open file: $!";
# print "Storing Snyderome variant data\n";
# while(my $line = <SVAR>){
# 	next if $line =~ m/^#/;
# 	my ($chr, $coord) = ( split(/\t/, $line) )[0,1];
# 	$var_h{ $chr }{ $coord } = 1;
# }

#  dbsnp variants
# open(DBSNP, "All_20170403.vcf") || die "Can't open file: $!";
# print "Storing dbSNP variant data\n";
# while(my $line = <DBSNP>){
# 	next if $line =~ m/^#/;
# 	my ($chr, $coord) = ( split(/\t/, $line) )[0,1];
# 	$chr = 'chr'.$chr;
# 	$var_h{ $chr }{ $coord } = 1;
# }

# Iterate through each file and tally the number of discrepant/non-discrepant loci
# Also keep track of read-depth at each locus

my %record_h;
my $chr = $1 if $file_list[0] =~ m/(chr.*)_SRR.*\.rpileup/;

foreach my $file ( @file_list ){
	open (PILEUP, "$file") || die "Can't open file: $!";

	while (my $line = <PILEUP>){
		chomp ($line);
		my ($chr, $bp, $strand, $nuc, $depth) = split(/\t/, $line);

		next if exists $var_h{ $chr }{ $bp };
		if (!exists $record_h{ $bp }{ $nuc }){
			$record_h{ $bp }{ $nuc } = 0;
		}
		$record_h{ $bp }{ $nuc } += $depth;
	}
}

my $outfile = $chr.'_summary.txt';
open (OUT1, ">$outfile") || die "Can't open file: $!";

foreach my $bp (sort {$a<=>$b} keys %record_h) {
	my $comb_depth = 0;
	my @temp_array = ();
	foreach my $nuc ( ('A', 'C', 'G', 'T', '-') ){
		my $depth = 0;
		$depth = $record_h{$bp}{$nuc} if exists $record_h{$bp}{$nuc};
		$comb_depth += $depth;
		push @temp_array, $depth;
	}
	next if $comb_depth < 500;
	print OUT1 $chr, "\t", $bp, "\t", join("\t", @temp_array), "\n";
}
```
