## Supplementary material for "Antibody cross-reactivity accounts for widespread appearance of m^1^A in 5’ UTRs": code: download_fastq.sh

```
#!/bin/bash -l
#$ -j y
#$ -cwd
#$ -l os=rhel6.3
#$ -l zenodotus=true
#$ -l h_vmem=4G
#$ -N dwnld_fastq

# Store command line variable
sraid=`sed -n "${SGE_TASK_ID}p" $sraid_list `
echo "$sraid"

# Download data
cd $TMDPIR
~/bin/sratoolkit.2.8.0-ubuntu64/bin/fastq-dump $sraid
mv SRR* $SEQ/primary_data/rna_seq/snyder_GSE33029/smallRNA
```
