## Supplementary material for "Antibody cross-reactivity accounts for widespread appearance of m^1^A in 5’ UTRs": code: filter_variants_and_edit_sites.pl

```
#!/usr/bin/perl
use strict;
use warnings;

my $file = $ARGV[0];

# Snyder called variants
open(SVAR, "variants/TB0001907.all.ILLUMNIA.bwa.CEU.high_coverage.20101118.snp.raw.vcf") || die "Can't open file: $!";
my %var_h;
print "Storing Snyderome variant data\n";
while(my $line = <SVAR>){
	next if $line =~ m/^#/;
	my ($chr, $coord) = ( split(/\t/, $line) )[0,1];
	$var_h{ $chr }{ $coord } = 1;
}

#  dbsnp variants
open(DBSNP, "variants/All_20170403.vcf") || die "Can't open file: $!";
print "Storing dbSNP variant data\n";
while(my $line = <DBSNP>){
	next if $line =~ m/^#/;
	my ($chr, $coord) = ( split(/\t/, $line) )[0,1];
	$chr = 'chr'.$chr;
	$var_h{ $chr }{ $coord } = 1;
}

open(PILEUPSUMMARY, "$file") || die "Can't open file: $!";
open(OUT, ">output.txt") || die "Can't open file: $!";

print "Searching pileup summary for variants\n";
while(my $line = <PILEUPSUMMARY>){
	my ($chr, $coord) = ( split(/\t/, $line) )[0,1];
	if (! exists $var_h{ $chr }{ $coord }){
		print OUT $line;
	}
	
}
```
