## Supplementary material for "Antibody cross-reactivity accounts for widespread appearance of m^1^A in 5’ UTRs": code: final_rpileup_analysis.R

```
library(data.table)

# Read in pileup summary data for all chromosomes
pileup.summary <- fread("data/all_summary.txt", 
                        col.names=c('chr', 'end', 'a', 'c', 'g', 't', 'del'))

# Read in annotation data file from MetaPlotR (See GitHub page) that contains the gene information for every nucleotide
genic.annot <- fread("data/hg19_annot.sorted.bed",
                     col.names=c('chr', 'start', 'end', 'desc', 'nuc', 'strand'))

uniq.genic.annot <- genic.annot[!duplicated(genic.annot[,c("chr", "start", "end")]),]

pileup.summary.annot <- merge(pileup.summary, uniq.genic.annot, by = c("chr", "end"))
pileup.summary.annot$start <- rowSums(pileup.summary.annot[,c('a', 'c', 'g', 't')])
colnames(pileup.summary.annot)[8] <- "depth"

# Limit sites to those with read depth > 500
pileup.summary.annot <- pileup.summary.annot[pileup.summary.annot$depth > 500,]


# identify candidate modification sites
# select sites where mismatch rate is >= 5%
temp <- pileup.summary.annot[,c('a', 'c', 'g', 't')]/pileup.summary.annot$depth
pileup.summary.annot$alt.freq <- round(1-apply(temp,1,max), 2)

hits.p05 <- subset(pileup.summary.annot, alt.freq > 0.05)
output <- data.frame( hits.p05[,c('chr', 'end')], rep('', length(hits.p05$chr)))

# print out list of candidate hits
#write.table(output, "data/candidate_mods.txt", sep = "\t", col.names = FALSE, 
#            row.names = FALSE, quote = FALSE)

# Remove likely SNVs by picking only sites where the second most common nucleotide has a frequency of at least 1%

temp <- rep(FALSE, dim(nuc.frac)[1])
for (i in 1:dim(nuc.frac)[1]){
  nuc.list <- sort(nuc.frac[i,])
  if(nuc.list[2] > cutoff){
    temp[i] <- TRUE
  }
}
summary(temp)

#write.table(hits.p05[temp,], "data/final_pileup_hits.txt", row.names = F, quote=F, col.names=F, sep="\t")
```
