## Supplementary material for "Antibody cross-reactivity accounts for widespread appearance of m^1^A in 5’ UTRs": code: index_hg19_STAR.sh

```
#!/bin/bash -l
#$ -j y
#$ -cwd
#$ -l os=rhel6.3
#$ -l zenodotus=true
#$ -l h_vmem=6G
#$ -pe smp 16
#$ -N index_STAR

# Make directory for input and output
echo 'Create directories for input and ouput'
cd $TMPDIR
mkdir indexed

# Transfer data to local scratch
echo 'Transfer data to local scratch'
#rsync -avh ${SCRATCH}/Projects/seq_mods/primary_data/rna_seq/snyder_GSE33029/${sraid}* ${TMPDIR}/fastq/
rsync -avh $SCRATCH/genomes/human/ucsc/ucsc.hg19.fasta ./
rsync -avh $SCRATCH/annotations/human/hg19_ucsc_genes_annot.gtf ./

# Issue command
echo 'Running STAR'
$HOME/bin/STAR-STAR_2.4.2a/bin/Linux_x86_64/STAR \
    --runThreadN $NSLOTS \
    --runMode genomeGenerate \
    --genomeDir indexed \
 	--genomeFastaFiles ucsc.hg19.fasta \
    --sjdbGTFfile hg19_ucsc_genes_annot.gtf \
    --sjdbOverhang 100

# Transfer output back to scratch
rsync -avh indexed $SCRATCH/indexes/star/101bp_ucsc_hg19
```
