## Supplementary material for "Antibody cross-reactivity accounts for widespread appearance of m^1^A in 5’ UTRs": code: index_rRNA_STAR.sh

```
#!/bin/bash -l
#$ -j y
#$ -cwd
#$ -l os=rhel6.3
#$ -l zenodotus=true
#$ -l h_vmem=1G
#$ -pe smp 4
#$ -N index_rRNA

# Make directory for input and output
echo 'Create directories for input and ouput'
cd $TMPDIR
mkdir fasta
mkdir index

# Transfer data to local scratch
echo 'Transfer data to local scratch'
rsync -avh ${SCRATCH}/genomes/human/rRNA.fasta ${TMPDIR}/fasta/

# Issue command
echo 'Running STAR: indexing genome'

$HOME/bin/STAR-STAR_2.4.2a/bin/Linux_x86_64/STAR \
	--runThreadN $NSLOTS \
	--runMode genomeGenerate \
	--genomeDir $TMPDIR/index \
    --genomeFastaFiles $TMPDIR/fasta/rRNA.fasta \
    --genomeSAindexNbases 4


# Transfer output back to scratch
rsync -avh ${TMPDIR}/index/ $SCRATCH/indexes/star/101bp_hsa_rRNA/
```
