## Supplementary material for "Antibody cross-reactivity accounts for widespread appearance of m^1^A in 5’ UTRs": code: README.txt

Download Snyderome fastq files from GEO:
- download_fastq.sh

Index human genome (hg19) and rRNA with STAR:
- index_rRNA_STAR.sh
- index_hg19_STAR.sh

Align Snyderome fastq files (20 replicates) to the human genome and rRNA:
- rRNA_snyder_STAR.sh
- trx_snyder_STAR.sh

Run samtools Rpileup on each fastq file (shell script to run R script)
- Rpileup.sh
- rpileup.R

Split each pileup file by chromosome (shell script to run perl script)
- run_split_rpileup.sh
- split_rpileup_by_chr.pl

Summarize the results of the split pileup files.  Generate a single summary per chromosome containing the results from all 20 replicates. shell script to run perl script) 
- run_analyze_split_rpileup_method2.sh
- analyze_split_rpileup_method2.pl 

Filter results for genomic variants identified in dbSNP or from the Snyderome   genomic vcf
- run_filter_variants.sh
- filter_variants_and_edit_sites.pl

Combine the summary files separated by chromosome into one summary file containing all chromosomes (command line):
- cat *_summary.txt > all_summary.txt

Some basic cleanup/analysis of the summary file producing a final table of candidate modification sites
- final_rpileup_analysis.R


chrom_list_major.txt
