## Supplementary material for "Antibody cross-reactivity accounts for widespread appearance of m^1^A in 5’ UTRs": code: rpileup.R

```
args <- commandArgs(trailingOnly=TRUE)

library(Rsamtools)
bf <- BamFile(args[1])

# Specify parameter constructor
p_param <- PileupParam(max_depth=1000000, min_mapq=0,min_base_quality=20)

# Run pileup
pileup.res <- pileup(bf, pileupParam = p_param)

# Print out
write.table(x = pileup.res, file = "output.txt", quote = FALSE, sep = "\t", 
            row.names = F, col.names = F)
```
