## Supplementary material for "Antibody cross-reactivity accounts for widespread appearance of m^1^A in 5’ UTRs": code: Rpileup.sh

```
#!/bin/bash -l
#$ -j y
#$ -cwd
#$ -l os=rhel6.3
#$ -l zenodotus=true
#$ -l h_vmem=32G
#$ -N rpileup	

# Store command line variable
sraid=`sed -n "${SGE_TASK_ID}p" $sraid_list `
echo "$sraid"

# Make directory for input and output
echo 'Create directories for input and ouput'
cd $TMPDIR

# Transfer data to temp directory
rsync -avh $SCRATCH/Projects/seq_mods/alignments2/snyder_trx_star/${sraid}'_Aligned.sortedByCoord.out.bam' ./
rsync -avh $SCRATCH/Projects/seq_mods/scripts/rpileup.R ./

# Run pileup
Rscript --verbose rpileup.R ${sraid}'_Aligned.sortedByCoord.out.bam'

# Transfer output back to scratch
echo 'Transfer data from temp to scratch'
rsync -avh output.txt $SCRATCH/Projects/seq_mods/rpileup/${sraid}.rpileup
```
