## Supplementary material for "Antibody cross-reactivity accounts for widespread appearance of m^1^A in 5’ UTRs": code: run_analyze_split_rpileup_method2.sh

```
#!/bin/bash -l
#$ -j y
#$ -cwd
#$ -l os=rhel6.3
#$ -l zenodotus=true
#$ -l h_vmem=96G
#$ -N analyze_rpile	

# Store command line variable
chr=`sed -n "${SGE_TASK_ID}p" $chr_list `
echo "$chr"

# Make directory for input and output
echo 'Create directories for input and ouput'
cd $TMPDIR

# Transfer data to temp directory  
rsync -avh $SCRATCH/Projects/seq_mods/rpileup_split/${chr}'_'* ./
rsync -avh $SCRATCH/Projects/seq_mods/variants/TB0001907.all.ILLUMNIA.bwa.CEU.high_coverage.20101118.snp.raw.vcf ./
rsync -avh $SCRATCH/Projects/seq_mods/scripts/analyze_split_rpileup_method2.pl ./

# Run perl script
perl analyze_split_rpileup_method2.pl ${chr}*

# Transfer output from temp dir to $SCRATCH
rsync -avh *summary.txt $SCRATCH/Projects/seq_mods/rpileup_split_summary
```
