## Supplementary material for "Antibody cross-reactivity accounts for widespread appearance of m^1^A in 5’ UTRs": code: run_filter_variants.sh

```
#!/bin/bash -l
#$ -j y
#$ -cwd
#$ -l os=rhel6.3
#$ -l zenodotus=true
#$ -l h_vmem=16G
#$ -N rem_variants	

# Store command line variable
chr=`sed -n "${SGE_TASK_ID}p" $chr_list `
echo "$chr"

# Make directory for input and output
echo 'Create directories for input and ouput'
cd $TMPDIR

# Transfer data to temp directory  
rsync -avh $SCRATCH/Projects/seq_mods/rpileup_split_summary/${chr}'_'* ./
rsync -avh $SCRATCH/Projects/seq_mods/scripts/filter_variants_and_edit_sites.pl ./

# Run perl script
perl filter_variants_and_edits_sites.pl ${chr}'_'*

# Transfer output from temp dir to $SCRATCH
rsync -avh output.txt $SCRATCH/Projects/seq_mods/rpileup_split_summary_filtered/${chr}'_rpileup_nuc_discrepancy_summary.txt'
```
