## Supplementary material for "Antibody cross-reactivity accounts for widespread appearance of m^1^A in 5’ UTRs": code: run_split_rpileup.sh

```
#!/bin/bash -l
#$ -j y
#$ -cwd
#$ -l os=rhel6.3
#$ -l zenodotus=true
#$ -l h_vmem=2G
#$ -N split_rpileup	

# Store command line variable
sraid=`sed -n "${SGE_TASK_ID}p" $sraid_list `
echo "$sraid"

# Make directory for input and output
echo 'Create directories for input and ouput'
cd $TMPDIR

# Transfer data to temp directory  
rsync -avh $SCRATCH/Projects/seq_mods/rpileup/${sraid}'.rpileup' ./
rsync -avh $SCRATCH/Projects/seq_mods/data/chrom_list_major.txt ./
rsync -avh $SCRATCH/Projects/seq_mods/scripts/split_rpileup_by_chr.pl ./

# Run perl script
perl split_rpileup_by_chr.pl ${sraid}*

rsync -avh ${chr}* $SCRATCH/Projects/seq_mods/rpileup_split
```
