## Supplementary material for "Antibody cross-reactivity accounts for widespread appearance of m^1^A in 5’ UTRs": code: split_rpileup_by_chr.pl

```
#!/usr/bin/perl
use strict;
use warnings;

my $file = $ARGV[0];
my $sraid = $1 if $file =~ m/(SRR.*)\.rpileup/;

## Store a list of the major chromosomes #######################
open (MAJCHR, "chrom_list_major.txt") || die "Can't open directory: $!";
my %major_chr;
while (my $chr = <MAJCHR>){
	chomp ($chr);
	$major_chr{$chr} = 1;
}

open (PILEUP, "$file") || die "Can't open file: $!";

my $previous_chr = '';
my $first_line = 'F';

while (<PILEUP>){
	my ($chr, $bp, $strand, $nuc, $depth) = split(/\t/, $_);
	next if !exists $major_chr {$chr}; 
	if ($chr ne $previous_chr and $first_line ne 'T'){
		close(OUT);
		my $outfile = $chr.'_'.$sraid.'.rpileup';
		open (OUT, ">$outfile") || die "Can't open file: $!";
		$previous_chr = $chr;
	}
	print OUT $_;
	$first_line = 'F';
}
```
