## Supplementary material for "Antibody cross-reactivity accounts for widespread appearance of m^1^A in 5’ UTRs": code: trx_snyder_STAR.sh

```
#!/bin/bash -l
#$ -j y
#$ -cwd
#$ -l os=rhel6.3
#$ -l zenodotus=true
#$ -l h_vmem=4G
#$ -pe smp 16
#$ -N trx_STAR

# Store command line variable
sraid=`sed -n "${SGE_TASK_ID}p" $sraid_list `
echo "$sraid"

# Make directory for input and output
echo 'Create directories for input and ouput'
cd $TMPDIR
mkdir fastq
mkdir bamout

# Transfer data to local scratch
echo 'Transfer data to local scratch'
rsync -avh $SCRATCH/Projects/seq_mods/primary_data/rna_seq/snyder_GSE33029/${sraid}* fastq/
rsync -avh $SCRATCH/indexes/star/101bp_ucsc_hg19/indexed ./

# Issue command
echo 'Running STAR'
$HOME/bin/STAR-STAR_2.4.2a/bin/Linux_x86_64/STAR \
    --runThreadN $NSLOTS \
    --genomeDir indexed \
    --readFilesCommand zcat \
    --readFilesIn fastq/${sraid}_1.fastq.gz fastq/${sraid}_2.fastq.gz \
    --outSAMattributes All \
    --outFilterType BySJout \
    --outFilterMultimapNmax 1 \
    --outFilterMismatchNmax 1 \
    --limitBAMsortRAM 4000000000 \
    --outSAMtype BAM SortedByCoordinate \
    --outFilterScoreMinOverLread 0 \
	--bamRemoveDuplicatesType UniqueIdentical \
	--clip3pNbases 10 \
	--clip5pNbases 10 \
    --outSAMattrRGline ID:id_${sraid}  PL:"ILLUMINA"   PU:"CXX_1"  SM:sample_${sraid}   LB:lib_${sraid} \
    --outFileNamePrefix bamout/${sraid}_

# Transfer output back to scratch
echo 'Transfer data from temp to scratch'
rsync -avh ${TMPDIR}/bamout/${sraid}* $SCRATCH/Projects/seq_mods/alignments2/snyder_trx_star
```
